# EGR1 transcriptional control of human cytomegalovirus latency

**DOI:** 10.1101/648543

**Authors:** Jason Buehler, Ethan Carpenter, Sebastian Zeltzer, Suzu Igarashi, Michael Rak, Iliyana Mikell, Jay A. Nelson, Felicia Goodrum

## Abstract

Sustained phosphotinositide3-kinase (PI3K) signaling is critical to the maintenance of herpesvirus latency. We have previously shown that the beta-herpesvirus, human cytomegalovirus (CMV), regulates epidermal growth factor receptor (EGFR), upstream of PI3K, to control states of latency and reactivation. Inhibition of EGFR signaling enhances CMV reactivation from latency and increases viral replication, but the mechanisms by which EGFR impacts replication and latency is not known. We demonstrate that HCMV downregulates MEK/ERK and AKT phosphorylation, but not STAT3 or PLCγ for productive replication. Similarly, inhibition of either MEK/ERK or PI3K/AKT, but not STAT or PLCγ, pathways increases viral reactivation from latently infected CD34^+^ hematopoietic progenitor cells (HPCs), defining a role for these pathways in latency. We hypothesized that CMV modulation of EGFR signaling might impact viral transcription. Indeed, EGF-stimulation increased expression of the *UL138* latency gene, but not immediate early or early viral genes, suggesting that EGFR signaling promotes latent gene expression. The early growth response-1 (EGR1) transcription factor is induced downstream of EGFR signaling through both PI3K/AKT and MEK/ERK pathways. EGR1 expression is important for the maintenance of HPC stemness and its downregulation drives HPC differentiation and mobilization. We demonstrate that EGR1 binds upstream of *UL138* and is sufficient to promote *UL138* expression. Further, disruption of EGR1 binding upstream of *UL138* prevented CMV from establishing a latent infection in CD34^+^ HPCs. Our results indicate a model whereby UL138 modulation of EGFR signaling feeds back to promote UL138 expression and suppression of replication to establish or maintain viral quiescence.

**AUTHOR SUMMARY:** CMV regulates EGFR signaling to balance states of viral latency and replication. CMV blocks downstream PI3K/AKT and MEK/ERK signaling pathways through down-regulation of EGFR at the plasma membrane. PI3K/AKT and MEK/ERK signaling increases expression of the EGR1 transcription factor that is necessary for the maintenance of stem cell stemness. A decrease in EGR1 expression promotes HPC mobilization to the periphery and differentiation, a known stimulus for CMV reactivation. We identified functional EGR1 binding sites upstream of the *UL138* gene and EGR-1 binding stimulates *UL138* expression. Additionally, down-regulation of EGR1 by CMV miR-US22 decreases *UL138* expression emphasizing the importance of this transcription factor in expression of this latency gene. Lastly, we demonstrate that a CMV mutant virus lacking an upstream EGR1 binding site is unable to establish latency in CD34^+^ HPCs. This study defines one mechanism by which EGFR signaling impacts viral gene expression to promote CMV latency.

## INTRODUCTION

The mechanisms by which herpesviruses persist through the establishment of a quiescent infection, known as latency, and reactivate for continued transmission are incompletely defined. It is known that herpesviruses sense and respond to changes in the host cell signaling, such as that associated with stress and differentiation, to modulate the decisions to maintain latency or to reactivate. However, the molecular underpinnings of how these cellular signals induce changes in chromatin and viral gene expression are less well defined. Human cytomegalovirus (CMV) is a beta-herpesvirus that persists within the majority of the human population. During infection of an immunocompetent host, CMV has a protracted acute phase and then establishes a life-latent infection, which is marked by sporadic subclinical reactivation events. CMV establishes latency in CD34+ hematopoietic progenitor cells (HPCs) and is carried through differentiation in cells of the myeloid lineage, including CD14+ monocytes (1). During latency in experimental models, CMV genes are expressed broadly but at very low levels (2, 3) and replication is restricted. Reactivation in immunodeficient individuals, such as stem cell or solid organ transplant recipients, is a major cause for morbidity and mortality (4–6). Additionally, CMV reactivation in patients undergoing intensive chemotherapy treatments can cause severe pathologies, including pneumonia, enteritis, blindness, and deafness(7, 8). Currently, there is no vaccine and existing antivirals have toxicity issues and cannot target latently infected cells. Understanding the molecular mechanisms that define and control the latent CMV infection is critical for the development of novel strategies to target the latent infection.

Virus manipulation of host cell signaling during infection of hematopoietic cells provides the means by CMV ensures survival of the infected cells and control differentiation and reactivation (9–14). Epidermal growth factor receptor (EGFR) signaling is a key component of the molecular switch regulating the establishment of latency and reactivation of viral replication (15). In CD34+ HPCs, CMV stimulates EGFR during entry and these initial signaling events are important for the establishment of latency (16). EGFR also serves as an entry receptor for CMV into fibroblasts (17), although sustained EGFR signaling represses productive replication (15).

The CMV UL135 and UL138 gene products antagonistize on another in regulating latency and reactivation (18). UL138 is suppressive to virus replication and critical for the establishment of latency, whereas UL135 is important for reactivation. UL135 functions, in part, by overcoming the suppressive effects of UL138, which otherwise block the initiation of viral replication from infectious genomes Accordingly, UL135 and UL138 gene products both interact with EGFR, but have opposing effects on the regulation of EGFR trafficking and signaling (15). UL135, reduces total and cell surface levels of EGFR. UL135 regulates EGFR trafficking and signaling through its interactions with the host adapter proteins for the Cbl E3 ubiquitin ligase, Abelson interacting protein 1 (Abi1), CIN85 and CD2AP (15, 19). Mutations in *UL135* ablating these host interactions restore EGFR levels in infected fibroblasts and diminish reactivation from latent infection (19). The mechanism by which UL138 regulates EGFR is less clear. However, in contrast to UL135, UL138 increases cell surface levels of EGFR in productively infected fibroblasts and is important for sustained EGFR signaling activity (15). The interactions between UL135 and Abi-1 and CIN85/CD2AP are required for reactivation, directly linking UL135- mediated degradation of EGFR to reactivation (20).

In this study, we defined the PI3K/AKT and MEK/ERK pathways downstream of EGFR as important to the establishment of latency using our experimental CD34+ HPC model. By contrast, the PLCγ and STAT pathways did not impact latency in our system. Additionally, EGF-stimulation increased UL138 gene expression, suggesting that EGFR signaling impacts latency gene expression. We mapped consensus binding sites for the early growth response factor 1 (EGR1) transcription factor upstream of *UL138.* EGR1 is highly expressed in CD34+ HPCs and is required to maintain stemness (21). Here, we show that EGR1 binds to sites within the *UL133-UL138* gene locus and stimulates UL138 gene expression and these sites are required to maintain latency in CD34^+^ HPCs. From these findings a positive feedback model emerges where by UL138 sustains EGFR signaling and EGFR signaling stimulates EGR1, which then drives UL138 gene expression. Disruption of EGR-1 regulation of UL138 expression results in a loss of latency and the virus replicates. This work defines one mechanism by which EGFR signaling impacts viral gene expression to regulate CMV latency and reactivation.

## RESULTS

### CMV downregulates total and cell surface levels of EGFR

We previously demonstrated that CMV modulates EGFR total and cell surface levels during infection in fibroblasts (productive infection) and CD34^+^ HPCs (site of latency)(15). In fibroblasts, we demonstrated that EGFR surface and total levels decrease substantially by 48 hours post infection (hpi). To further understand the regulation of EGFR during productive infection, we analyzed surface and total levels of EGFR in fibroblasts over a time course of infection from 0-72 hpi. Fibroblasts were infected with the TB40/E strain expressing GFP as a marker for infection (22), which serves as the parental/wild-type (WT) virus for all studies and EGFR surface levels were measured by flow cytometry (Fig. 1A). EGFR surface levels began to decrease by 12 hpi and were reduced to ~60% of uninfected cells by 24 hpi, and remained between 50 and 60% of uninfected cells for the remainder of the infection time course. Analysis of total EGFR levels over the same time course indicated that total EGFR levels were reduced to 40% and 20% of uninfected cells by 48 and 72 hpi, respectively (Fig. 1B). These findings are consistent with our previous work demonstrating the downregulation of EGFR during the productive cycle of infection (15, 20) and extend those observations by defining the onset of this downregulation as within the early stages of infection.

**Figure 1.**
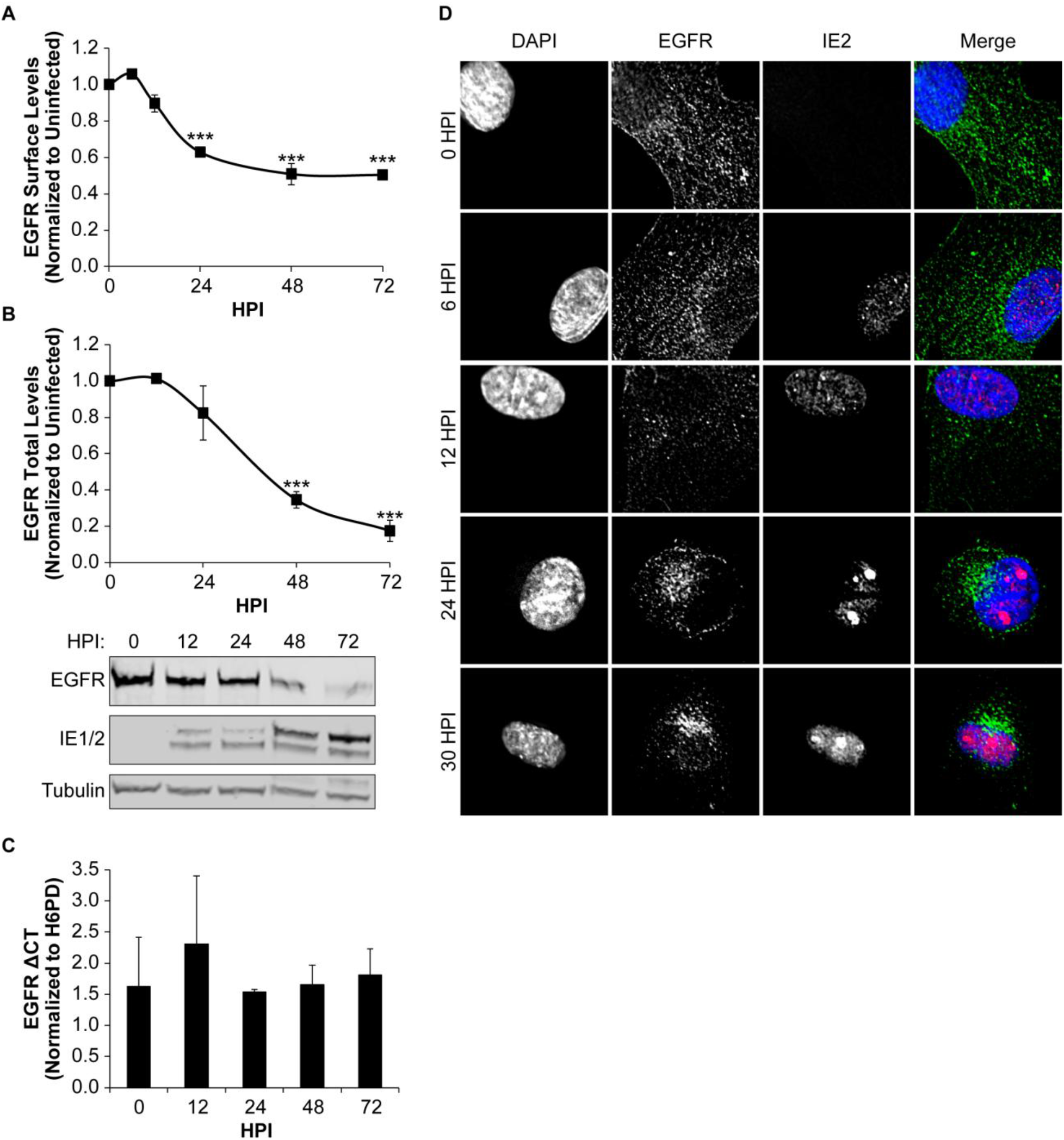
CMV downregulates EGFR surface and total protein levels as infection progresses. Fibroblasts were infected with TB40E_GFP_ virus at an MOI odf 1 for 0-72 hpi. (A) To measure EGFR surface levels, infected cells were stained with BV421 conjugated ms α-EGFR antibody and analyzed by flow cytometry. Normalized geometric mean fluorescent intensity is shown. (B) Total EGFR levels were measured over a time course by immunoblotting. Blots were stained with rb α-EGFR, ms α-IE1/2 antibody, and ms α-Tubulin. Both surface and total EGFR levels were normalized to 0 hpi for statistical analysis. IE proteins serve as a control for infection and tubulin serves as a control for loading. (C) Relative EGFR mRNA levels were measured over a time course using quantitative reverse transcriptase PCR and SYBR green. EGFR transcripts are normalized to H6PD, cellular housekeeping control, at each timepoint. (A-C) Statistical significance was calculated by One-Way ANOVA with Tukey’s correction and represented by asterisks (*** p-values < 0.001). Graphs represent the means from 3 independent replicates with error bars representing SEM. (D) Subcellular localization of EGFR was monitored over a time course of infection. Nuclei, EGFR and IE2 are visualized by staining with DAPI, rb α-EGFR, and ms α-IE2 and confocal deconvolution microscopy.

CMV was previously shown to transcriptionally downregulate EGFR (23, 24). While UL135 downregulates, total levels of EGFR (15, 20), disruption of UL135 or its interaction with host proteins does not fully restore EGFR to uninfected cell levels. This might be explained by a transcriptional downregulation of EGFR. However, we did not detect any significant alteration in EGFR mRNA levels by quantitative reverse transcriptase PCR (RT-qPCR) over the time course of infection(Fig. 1C). The lack of a transcriptional downregulation in our study may refelect difference in the virus strain used for infection or the cells type, and leaves open the possibility that other viral factors contribute to the diminishment of EGFR levels.

We previously observed a re-localization of EGFR to the viral assembly compartment during productive replication (15). To determine the timing of EGFR re-localization, EGFR subcellular localization was determined in TB40/E infected fibroblasts at (Fig. 1D). IE2 was used to mark infected cells. In uninfected cells (0 hpi) EGFR staining is localized at the plasma membrane and distributed in puncta throughout the cytoplasm. Although, the distribution of EGFR was not dramatically altered between 6 and 12 hpi, a large portion of EGFR re-localized at 24 hpi to a juxta-nuclear region and is maintained there. We previously demonstrated that this juxta-nuclear localization is proximal with markers, GM130 and pp28, for the viral assembly compartment (15). However, this result indicated that EGFR is re-localized at early times in infection prior to the formation of the assembly compartment.

### EGFR and downstream pathways are inhibited by CMV as replication progresses

To determine how CMV productive infection impacts EGFR signaling and pathways downstream of EGFR, we analyzed phosphorylation of EGFR, MEK1/2, ERK1/2, STAT3, PLCγ, and AKT at steady state or following 30 min of EGF stimulation in infected fibroblasts using a EGFR signaling array (PathScan, Cell Signaling Technologies). At steady state, phosphorylation of EGFR at T669, Y845, and Y1068 was increased relative to uninfected cells. However, EGFR was less responsive to EGF stimulation, marked by decreased phosphorylation on T669, Y845, and Y1068 relative to uninfected, stimulated cells (Fig. S1A). Infection did not alter phosphorylation of EGFR Y998. While basal activity of MEK1/2, ERK1/2, and AKT was not altered by infection in unstimulated cells, EGF-stimulated phosphorylation of MEK1/2 (S221 or S217/221, Fig. S1B) and AKT (S473, Fig. S1C) was reduced in infected cells relative to uninfected cells. CMV infection did not affect phosphorylation and activation of STAT3 (Y705), PLCγ1 (S1248) or AKT (T308) in response to EGF stimulation (Figure S1C). These results suggest that MEK/ERK and AKT signaling are the primary pathways downstream of EGFR that are suppressed by CMV infection.

To further analyze the inhibition of both AKT and MEK1/2 pathways by CMV, we monitored their activation over a time course of infection. Serum starved, infected fibroblasts were pulsed with EGF ligand over a time course following infection (0, 12, 24, 48, and 72 hpi) and cell lysates were harvested at 0, 15 and 30 minutes following each EGF pulse to analyze the phosphorylation of EGFR (Y1068), AKT (S473), and MEK1/2 (S217/221) (Fig. 2A). In uninfected fibroblasts, EGFR, AKT and MEK1/2 phosphorylation was induced by 15 min post EGF stimulation, as expected. In fibroblasts infected for 12 hours, pEGFR, pAKT, pMEK1/2 induction was unchanged from uninfected cells. In contrast, the levels of all three phosphorylation markers decreased significantly after EGF stimulation with a reduction of pMEK1/2 by 24 hpi and both pEGFR and pAKT by 48 hpi, relative to uninfected fibroblasts (Fig. 2B). By 72 hpi, pEGFR and pAKT was undectectable in infected cells (Fig. 2A). Induction of pMEK1/2 in response to EGF stimulation is undetectable relative to basal levels by 24 hpi. Together these data indicate that EGFR and both the MEK/ERK and PI3K/AKT downstream signaling nodes become progressively less responsive to stimulation in the productive infection.

**Figure 2.**
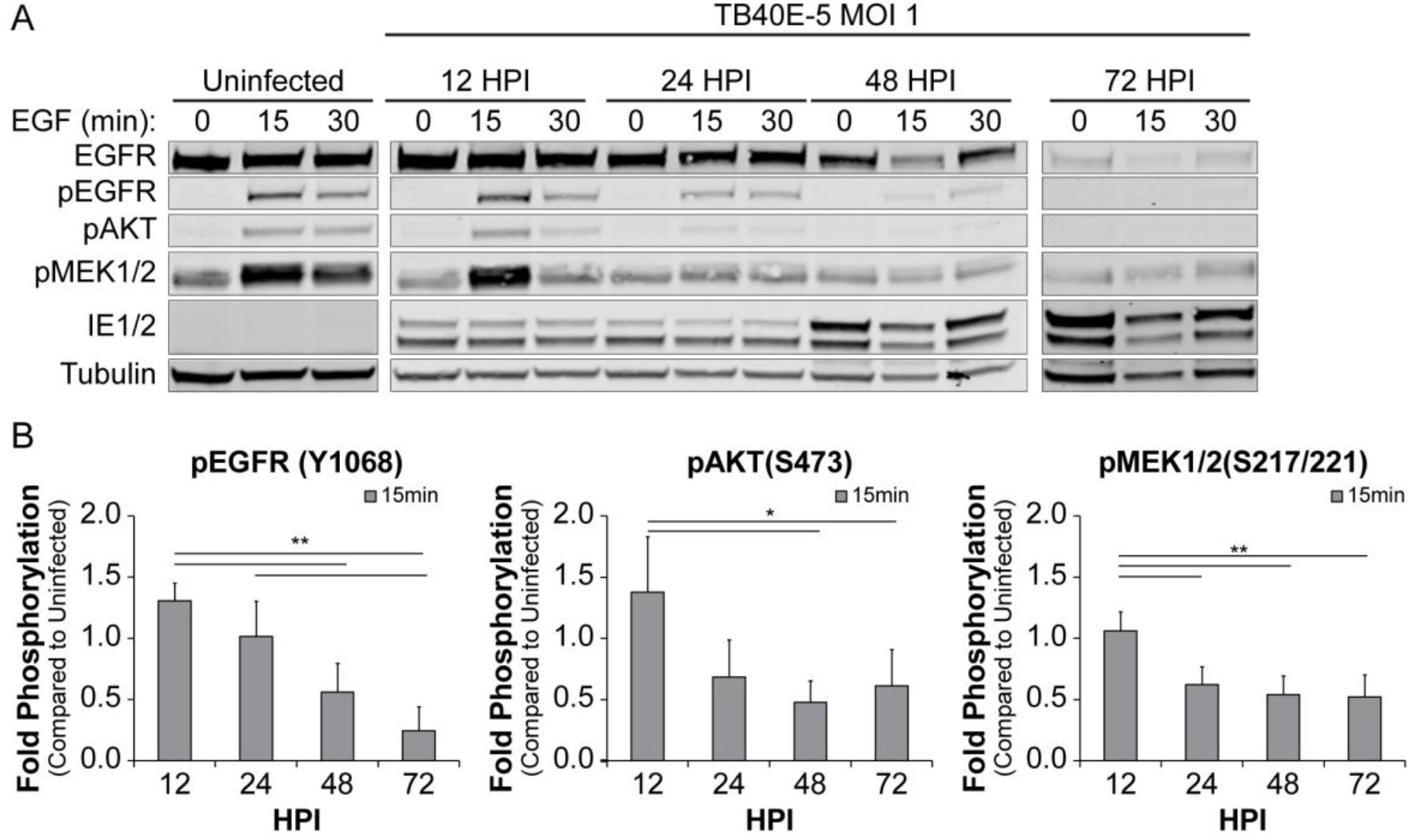
CMV infection prevents activation of AKT and MEK1/2. (A) Fibroblasts were serum starved for 24h and cells were then infected for 0-72 hpi. At each timepoint, infected cells were pulsed with 10 nM of EGFR for 0-30 min, and lysed. Lysates were separated out on SDS-PAGE gel, transferred on PVDF membrane, and stained for rb α-EGFR, rb α-pEGFR(Y1068), rb α-pAKT(S472), rb α-pMEK1/2(S217/221), ms α-IE1/2 antibody, and ms α-Tubulin. (B) The 15 min post EGF timepoint for all phosphorylation markers were normalized to uninfected cells and graphed to calculate statistics. Statistical significance was calculated by One-Way ANOVA with Tukey’s correction and represented by asterisks (* p-value < 0.05 and ** p-value < 0.01).Graphs represent the mean of three replicates and error bars represent SEM.

### PI3K/AKT and MEK/ERK pathways suppress viral replication for latency

Work by the Chan and Yurochko groups have demonstrated that PI3K signaling is important to survival of CMV-infected monocytes (10, 25). Further, we previously demonstrated that inhibition of EGFR or PI3K increases CMV replication in fibroblasts and reactivation in CD34^+^ hematopoietic progenitor cells (HPCs) (15). To further investigate how pathways downstream of EGFR impact CMV replication and latency, we used chemical inhibitors of the MEK/ERK, STAT, PI3K/AKT, and PLCγ pathways. The efficacy of each inhibitor at the chosen concentration was confirmed by analyzing phosphorylation over a 5 day time course (Fig. S2). Fibroblasts were treated with inhibitors at 1 day post infection so as not to interfere with viral entry (16, 26, 27) and viral titers were measured at 8 dpi (Fig. 3A). Inhibition of PI3K (LY294002) or AKT (MK-2206) increased viral titers by 7.6 and 7-fold, respectively, in comparison to the vehicle control, similar to what we have previously reported with EGFR inhibition (15). In contrast, inhibition of STAT1 (Fludarabine) or STAT3 (S3I-201) decreased virus production. Loss of viral replication by STAT inhibition has previously been reported with these and similar inhibitors (28, 29). The diminishment of virus replication was not related to cytotoxicity, as the monolayers stayed intact. Inhibition of MEK1/2 (Binimetinib), ERK1/2 (SCH772984), or PLCγ (U73122) did not alter virus titers relative to the vehicle control. These data confirm that PI3K/AKT pathways are suppressive to virus replication and demonstrate that MEK/ERK and PLCγ pathways are dispensable for productive infection in fibroblasts.

**Figure 3.**
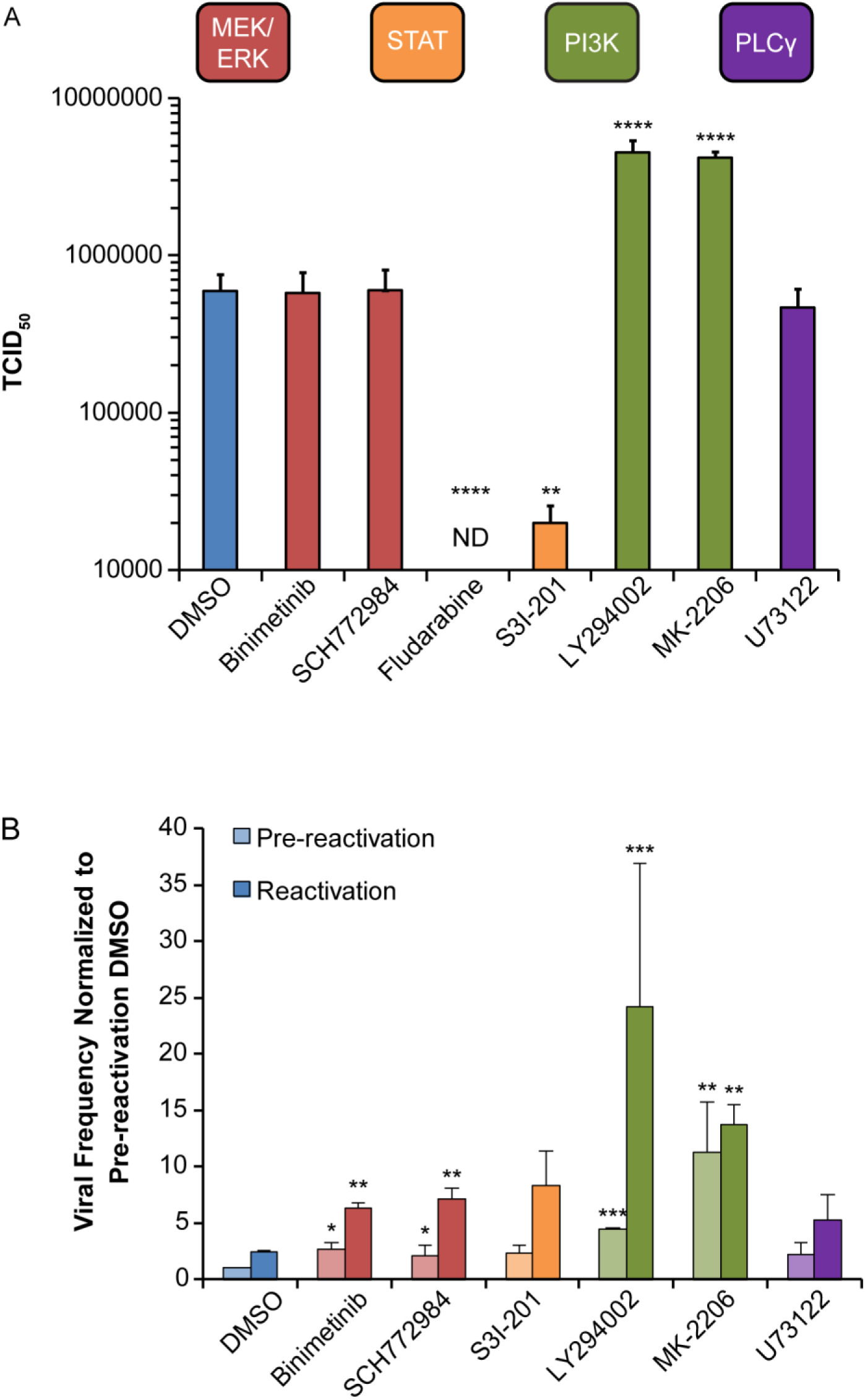
Inhibition of MEK/ERK and PI3K/AKT signaling stimulates reactivation in CD34^+^ HPCs, but only inhibition of PI3K/AKT stimulates replication in fibroblasts. (A) Fibroblasts were infected with TB40E_GFP_ virus (MOI=1). At 24 hpi, cells were treated with DMSO, MEK/ERK inhibitors (Binimetinib 1 μM; SCH772984 125nM), STAT (Fludarabine 50 μM; S3I-201 100 μM), PI3K/AKT (LY294002 20 μM; MK-2206 1.25 μM), PLCγ (U73122 4 μM). At 8 dpi media and cells were collected and viral titers were determined by TCID_50_. (B) CD34^+^ HPCs were infected with TB40E_GFP_ virus (MOI=2). At 24 hpi, CD34^+^/GPF^+^ cells were sorted and put into long-term culture with inhibitor list above. After 10 days, parallel populations of either mechanically lysed cells or whole cells were plated onto fibroblasts monolayers in cytokine-rich media and frequency of infectious centers was determined by limited dilution analysis. The mechanically lysed population defines the quantity of virus present prior to reactivation (pre-reactivation). The whole cell population undergoes differentiation due to fibroblasts contact and cytokine stimulation, which promotes viral reactivation (reactivation). The frequency was normalized to the pre-reactivation DMSO control to facilitate comparisons between experiments. Statistical significance was calculated by One-Way ANOVA with Tukey’s correction for each condition and represented by asterisks (* p-value < 0.05, ** p-value < 0.01, *** p-value < 0.001, and **** p-value < 0.0001). For fludarabine infected fibroblasts ANOVA could not measure difference due to absence of quantifiable virus and statistical significance was calculated by student t-test (**** p-value < 0.0001). Data graphed is the mean of 3 replicates with error bars representing SEM.

To determine the importance of the signaling pathways downstream of EGFR to latency and reactivation, we analyzed the impact of each inhibitor on latency and reactivation in CD34+ HPCs.. Infected (GFP+) CD34^+^ HPCs were isolated by fluorescent activated cell sorting (FACS) and co-cultured for 10 days in long-term bone marrow cultures using a bone marrow stromal cell support that has been shown to maintain hematopoietic cell progenitor phenotype and function (30). This period in long-term bone marrow culture allows for the establishment of CMV latency. At 10 dpi, half of the cells were seeded by limiting dilution into co-culture with fibroblasts in a cytokine-rich media to promote myeloid cell differentiation and reactivation. The other half of the culture was lysed and seeded by limiting dilution in parallel onto fibroblasts to quantify virus formed during the latency period (pre-reactivation) (31). Reactivation resulted in a 2-3 fold increase in the frequency of infectious centers relative to the pre-reactivation control (DMSO control, Fig. 3B). Inhibition of MEK/ERK with Binimetinib or SCH772984 increased the frequency of infectious centers by at least 2-fold in both the pre-reactivation and reactivation condition relative to their respective DMSO control. Inhibition of PI3K (LY294002) induced a 4-fold increase in infectious centers in the pre-reactivation and an almost 10-fold increase in infectious centers in the reactivation. Inhibition of AKT (MK-2206) resulted in a loss of latency with a 10-fold increase in the frequency of infectious centers in the pre-reactivation and a 5.5-fold increase in the reactivation, relative to DMSO controls. By contrast, inhibition of STAT3 and PLCγ did not significantly alter infectious centers produced prior to or following reactivation relative to the DMSO controls. Fludarbine was not used in CD34+ HPCs because a non-toxic dose could not be found. These results indicate that the MEK/ERK and PI3K/AKT pathways each contribute to the maintenance of CMV latency and inhibition of these pathways enhances reactivation, while we find no role for STAT3 or PLCγ.

### EGF stimulation drives expression of the *UL138* latency determinant

Collectively, our work demonstrates a requirement for EGFR and downstream PI3K/AKT and MEK1/2 signaling pathways for the suppression of virus replication to maintain latency in CD34+ (Fig. 2B) (15). However, the mechanisms by which host signaling impacts infection is not know. While the effect of EGFR signaling on cellular gene expression (32) might impact virus replication and latency, we hypothesized that EGFR signaling might also impact viral gene expression and, specifically, the expression of genes required for latency.

To determine if stimulation of EGFR might affect viral gene expression from the *UL133-UL138* locus, we monitored expression of the immediate early genes *UL122* and *UL123* (IE2 and IE1, respectively), *UL135*, and *UL138* in serum starved, infected fibroblasts over a time course following EGF stimulation (Fig. 4A). UL138 protein accumulation increased by 75% at 1h following EGF stimulation relative to unstimulated cells (Fig. 4A). UL138 protein levels remained elevated for up to 6h post EGF pulse. In contrast, neither *UL135* nor IE1/2 levels changed in response to EGF-stimulation.

**Figure 4.**
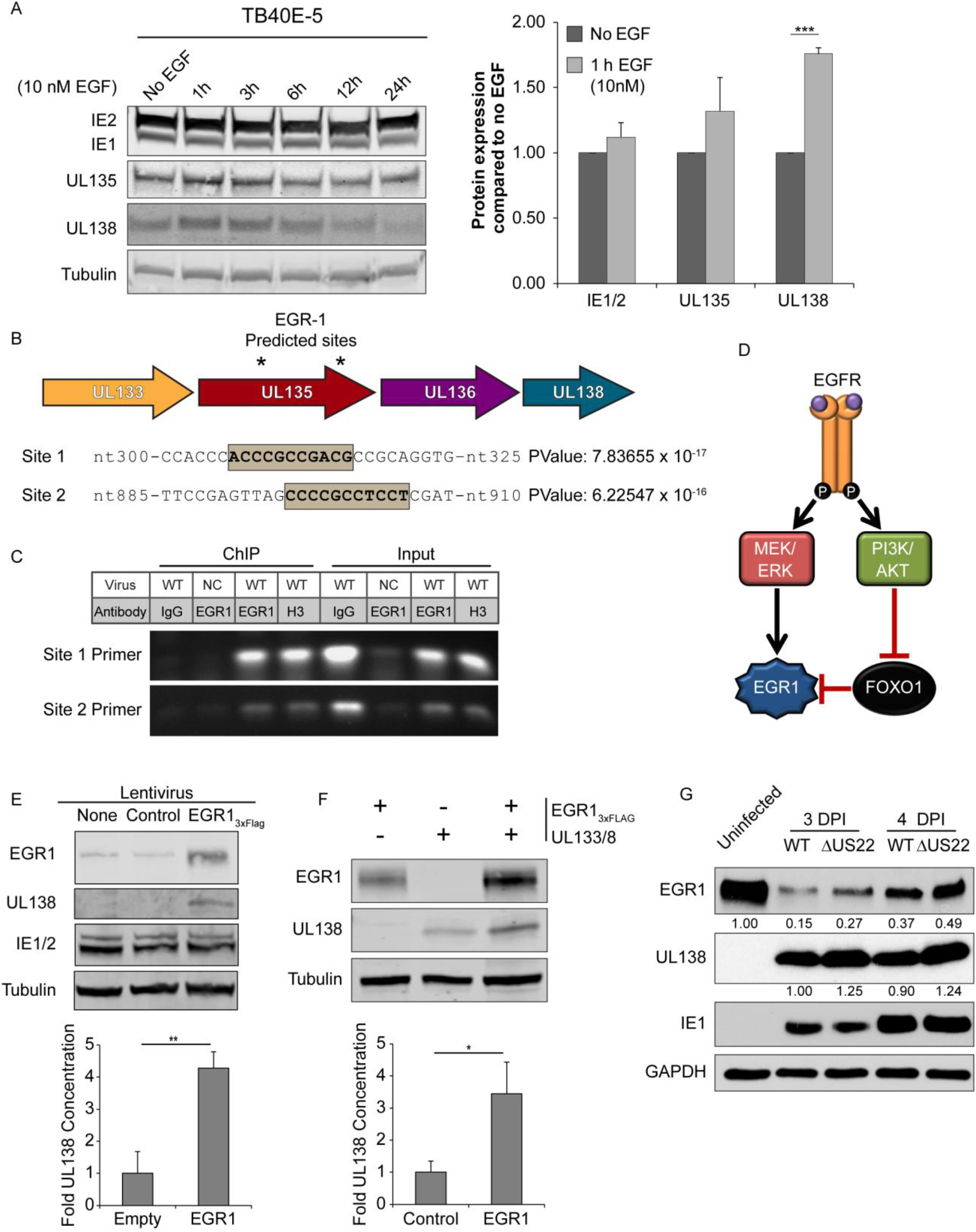
EGF-stimulation promotes *UL138* expression through EGR1 expression. (A) Fibroblasts were infected with TB40E_GFP_ (MOI=1) and put into serum-free media at 24hpi. Cells were stimulated with 10nM EGF at 48 hpo and cells were harvested between 1 and 24 hours post stimulation. Lysates were separated by SDS-PAGE, transfer onto a member, and blotted with ms α- IE1/2, rb α-*UL135*, rb α-*UL138*, and ms-α Tubulin. Protein levels from 4 replicates were normalized to no EGF treated control and 1h post EGF treatment is graphed. Statistical significance was calculated by student t-test; asterisks *** p-value < 0.001. Error bars represent SEM.(B) Graphical representation of putative EGR1 binding sites located within *UL135* ORF starting at nt-306 and nt-896, in reference to UL135 start codon. P-values were calculated using PhysBinder prediction software. (C) Fibroblasts were transduced with EGR1_3xFlag_ lentivirus and then infected with wildtype or *UL133/8*_*nul*l_ mutant (negative control; NC) TB40E virus (MOI=1). Chromatin was immunoprecipitated (ChIP) with IgG or antibodies specific to EGR1 or histone 3 (H3) and EGR1 binding Site 1 or Site 2 was detected in the precipitates by PCR. As a control, PCR was also performed on 2% of the ChIP input. Gel is a respresentative experiment from 3 replicates. (D) A model demonstrating how EGFR signaling promotes EGR1 expression by either directing its expression through MEK/ERK signaling or by blocking FOXO1 suppression of EGR1 transcription though PI3K/AKT signaling. (E) Fibroblasts were transduced with either EGR1_3xFlag_ or Empty vector control and after 24h infected with TB40E_GFP_. At 48 hpi, protein lysates were collected, separated by SDS-PAGE and blotted using rb α-FLAG, rb α-*UL138*, and ms α-Tubulin. (F) HEK293T cells were co-transfected with EGR1_3xFlag_, *UL133/8* encoding plasmid, or empty vector (minus sign). After 48h, samples were separated by SDS-PAGE and blotted for ms α-FLAG, rb α-*UL138*, and ms α-Tubulin. (E-F) *UL138* protein levels from either 4(E) or 3(F) independent experiments were normalized to control and shown in graphs. Statistical significance was calculated by student t-test; asterisks indicate * p-value < 0.05 and ** p-value < 0.01. Error bars represent SEM. (G) Fibroblasts were infected with 1 MOI of wild type or ΔmiR-US22 TB40E_GFP_ virus and serum starved overnight before treating with 50 ng/mL of EGF. Samples were collected at 3 and 4 dpi, then separated on a SDS-PAGE gel, and blotted for ms α-EGR1, rb α-*UL138*, ms α-IE1, and ms α-GAPDH. Normalized values for EGR1 and *UL138* protein are below each band. Blot is representative of two independent experiments.

To begin to understand how EGFR signaling might affect *UL138* expression, we used PhysBinder to identify putative transcription factor binding sites within the *UL133-UL138* locus that are regulated by EGFR signaling (33). To minimize the potential for false positives, we used the Max Precision setting and identified two binding sites for early growth response factor 1 (EGR1) within the *UL135* open reading frame (ORF) upstream of *UL138* (Fig. 4B). To confirm binding of EGR1 to these sites, we transduced fibroblasts with lentiviruses expressing EGR1 fused to a 3xFlag epitope tag (EGR1_3xFlag_) and infected the cells with either a wild-type or a UL133/8 deletion mutant (negative control; NC) TB40/E viruses. At 48 hpi, samples were processed for chromatin immunoprecipitation (SimpleChIP, Cell Signaling Technologies) with normal IgG or antibodies to either EGR1 or Histone 3 (H3) and binding was detected by PCR with site-specific primers (Fig. 4C). The EGR1 antibody precipitated sequences in the wild-type infection for both binding sites by PCR; but not in the NC or IgG control. These results indicate that EGR1 interacts with both site 1 and site 2. Interesting, both PI3K/AKT and MEK/ERK pathways induce EGR1 (34–36)(Fig. 4D). EGR-1 is of particular interest because it is highly expressed in hematopoietic cells, a site of CMV latency (21, 37). EGR1 is required to maintain stem cell quiescence and retention in the bone marrow and must be downregulated for hematopoietic differentiation and migration out of the bone marrow. Therefore, we hypothesized that EGR1 might stimulate the transcription of RNAs encoding *UL138* that we previously mapped to initiate downstream of the *UL135* ORF (38–40). The presence of putative EGR1 binding sites upstream of these transcripts suggests the existence of a promoter element in this region to regulate *UL138* expression; this element has not been mapped yet. In this case, EGR1 binding would be expected to induce *UL138*, but not *UL135* gene expression, consistent with our result in Figure 4A.

To determine if EGR1 is sufficient to induce *UL138* expression we transduced fibroblasts with lentivirus expressing either EGR1_3xFlag_ or empty vector and infected cells with TB40/E. EGR1 overexpression increased *UL138* expression in infected cells by 4-fold, while IE 1 and 2 proteins were unaffected (Fig. 4E). To ensure that EGR1 activity is not priming UL138 expression from a more distal promoter site, we co-transfected HEK-293T cells with a plasmid containing the entire *UL133-UL138* locus in a promoterless vector backbone (UL133/8) and either the EGR1_3xFlag_ expression construct or an empty control. UL138 can be expressed from this vector due to the presence of an IRES upstream of UL138 (38)however, UL138 protein levels increased 3.5-fold in cells overexpressing EGR1 relative to the empty vector control (Fig. 4F). These results indicate that EGR1 expression is sufficient to stimulate *UL138* expression from an unmapped regulatory element encoded with the *UL133/8* locus.

Mikell et al. have identified a CMV micro RNA transcribed within the US22 open reading frame (miR-US22) that targets EGR1, resulting in a 2 to 5-fold decrease in EGR1 protein levels depending on cell type and is required for reactivation (41). A miR-US22-mutant (ΔUS22) virus results in increased expression of EGR1 and fails to reactivate. We hypothesized ΔUS22 would also have increased expression of UL138. To test this, we infected fibroblasts with a wild type or ΔUS22-mutant TB40/E virus and pulsed them with EGF for 1 hour at different time points (Fig. 4G). At both 3 dpi and 4 dpi, UL138 was increased by ~25% in the ΔUS22-mutant virus infection relative to WT infection. The increase in UL138 protein levels corresponded to increased EGR1 protein levels in the context of ΔUS22 virus infection. This result indicates that UL138 gene expression is induced by EGFR signaling through the induction of EGR1 and suggests an epistatic relationship between UL138 and US22 and EGR1 in regulating infection.

### CMV maintains EGR1 levels during latent infection, but reduces its expression during productive replication

To determine how EGR1 is regulated during CMV latent infection, we analyzed EGR1 mRNA expression in TB40/E-infected CD34^+^ HPCs derived from two donors at 2 and 6 dpi by RNA sequencing (2). In each donor, EGR1 expression increased following CMV infection from 2 to 6 dpi by 3-fold(Fig. 5A). By contrast, the expression of two related genes belonging to the same zinc-finger transcription factor family, EGR2 and EGR3, were unchanged by CMV infection. Additionally, CMV did not affect expression of Wilms tumor 1 (WT1) a factor that binds competitively to the EGR1 consensus sequence to antagonize EGR1 transcriptional control of genes, including EGFR (42, 43). By contrast, EGR1 is suppressed 3-fold during replication in fibroblasts following an initial induction (Fig. 5B) that is likely due to the stimulation of EGFR at the cell surface during viral entry (16, 44). Further, EGR1 is strongly induced in serum starved, uninfected fibroblasts and infection diminishes this induction by 7-fold (Fig. 5C). The lack of responsiveness of EGR1 to EGF stimulation in infected cells likely reflects diminished EGFR levels and signaling in the context of infection beginning at 24 hpi (Figs. 1 and 2). These findings are consistent with the differential regulation of EGFR during productive and latent states of viral infection which we previously described (15).

**Figure 5.**
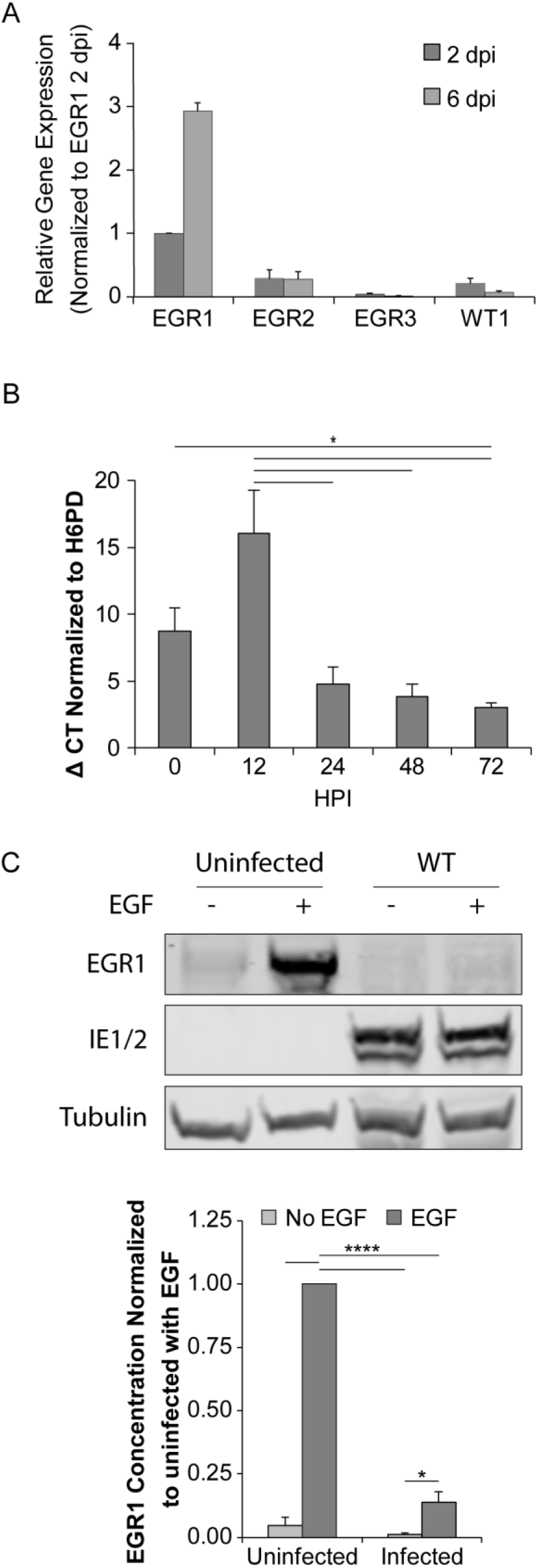
EGR1 levels are elevated during latent infection in CD34^+^ HPCs, but not during replication in fibroblasts. (A) CD34^+^ HPCs were infected with TB40E_GFP_ (MOI=2). At 2 and 6 dpi, we isolated RNA and prepared mRNA libraries for Illumina sequencing. Relative expression of EGR1, EGR2, EGR3, and WT1 was calculated by fragments per kilobase per million reads (FPKM) and normalized to EGR1 2 dpi levels. Error bars represent the range of gene expression between two independent donors. (B) Fibroblasts were infected with TB40E_GFP_ (MOI=1) and RNA was isolated at 0-72 hpi. EGR1 mRNA was quantified relative to H6PD by RT-qPCR. Results from 3 independent replicates are graphed error bars represent SEM. Statistical significance was calculated by One-Way ANOVA with Tukey’s correction and represented by an asterisk (* p-value < 0.05). (C) Fibroblasts were infected with TB40E_GFP_ (MOI=1) and transferred to serum-free media at 24 hpi. At 48 hpi, samples were pulsed with 10 nM of EGF for 1 h. Lystates were separated by SDS-PAGE and immunoblotted with rb α-EGR1, ms α-IE1/2, and ms α-tubulin. EGR1 protein levels were normalized to the uninfected sample stimulated with EGF and the mean from 3 independent replicates is graphed. Error bars represent SEM. We calculated statistical significance by two-way ANOVA with Tukey’s correction and represented significance by asterisks (* p-value < 0.05; **** p-value < 0.0001).

### EGR1 binding in the UL133-UL138 region induces UL138 protein accumulation

To validate the EGR1 binding sites upstream of *UL138*, we introduced silent mutations into wobble codons within site 1 or site 2 by site directed mutagenesis in the promoterless *UL133*/*8* vector to generate ΔSite 1 or ΔSite 2 constructs. HEK-293T cells were co-transfected with an empty vector or EGR1-Flag expression vector (EGR1_3xFLAG_) and either WT *UL133*/*8* or ΔSite 1 or ΔSite 2. UL138 protein levels in cells expressing EGR1_3xFlag_ were normalized to that in cells transfected with empty vector (Fig. 6A). EGR1 overexpression induced UL138 protein accumulation 4-fold from the WT UL133/8 construct; however mutation of either Site 1 or Site 2 resulted in 2-fold reduces levels of UL138 protein.

**Figure 6.**
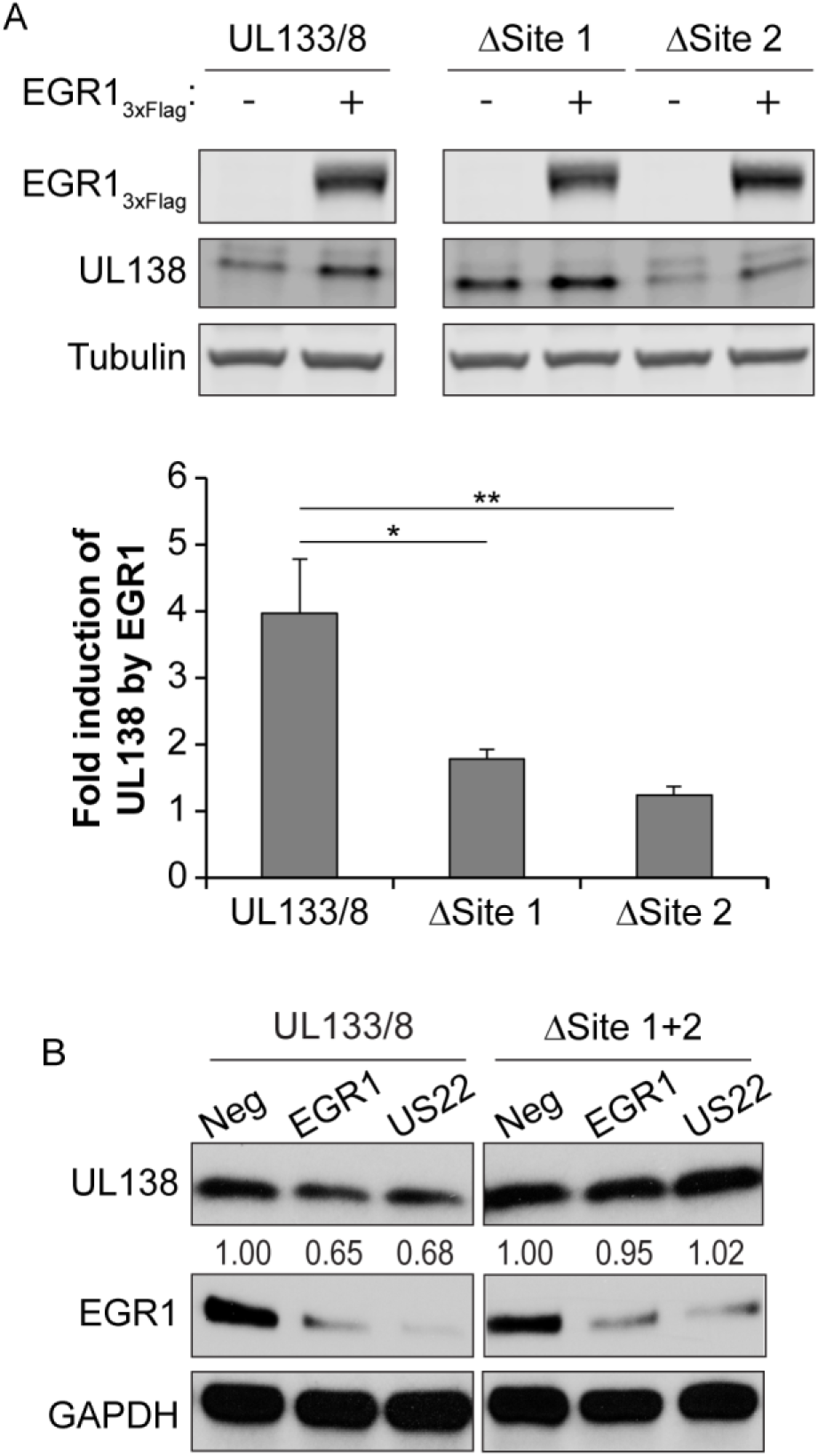
Mutation of EGR1 binding sites blocks induction of *UL138*. (A) HEK293T cells were co-transfected with either empty vector or EGR1_3xFlag_ and a plasmid contain UL133/8 or a mutant plasmid lacking one EGR1 binding site, ΔSite 1 or ΔSite 2. At 48 h lystates were separated by SDS-PAGE, and blotted for rb α-Flag, rb α-UL138, and ms α-tubulin. *UL138* protein levels in EGR1_3xFlag_ transfections were normalized to control levels to determine *UL138* induction. The results from 4 independent replicates are graphed. Statistical significance was calculated by One-Way ANOVA with Bonferroni correction (* p-value < 0.05 and ** p-value < 0.01). (B) HEK293T cells were cotransfected with the UL133/8 vector or the UL133/8 vector where EGR1 sites (ΔSite1, ΔSite 2) were disrupted and negative control siRNA, EGR1 siRNA, or miR-US22. Cells were transferred to serum-free media at 24 h. At 48 hpi, samples were stimulated with 50 ng/mL EGF for 1h and then lysed, separated by SDS-PAGE, and blotted for rb α-UL138 and ms α-GAPDH. *UL138* levels are normalized to negative control. A representative blot of 2 independent experiments is shown.

To further validate the role of the EGR-1 binding sites in regulating *UL138* expression, we analyzed UL138 protein levels in HEK 293T cells transfected with the WT UL133/8 or ΔSite1+2 promoterless vectors and EGR1 siRNAs or miR-US22 to knocked down EGR1 (Fig. 6B). Knockdown of EGR1 either with the EGR1 siRNA or miR-US22 decreased UL138 protein levels by 30% in cells containing the wild-type *UL133/8* plasmid. However, EGR1 knockdown had no effect on UL138 protein accumulation in cells where site 1 and site 2 was disrupted. Taken together, these results indicate the importance of EGR1 binding sites to EGR1-mediated induction of *UL138* expression.

We next engineered the EGR1 binding site disruptions into the TB40/E genome cloned as a bacterial artificial chromosome, resulting in TB40/E-ΔEGR1_Site 1_ and TB40/E-ΔEGR1_Site 2_. We confirmed the disruption of EGR1 binding sites by sequencing (Fig. S3). We evaluated each virus for its ability to replicate in fibroblasts by multi-step growth curves. Each virus replicated with wild-type kinetics and to wild type titers (Fig. 7A), indicating that the mutations introduced to disrupt EGR1 binding to the *UL133-UL138* region do not affect productive virus replication in fibroblasts.

**Figure 7.**
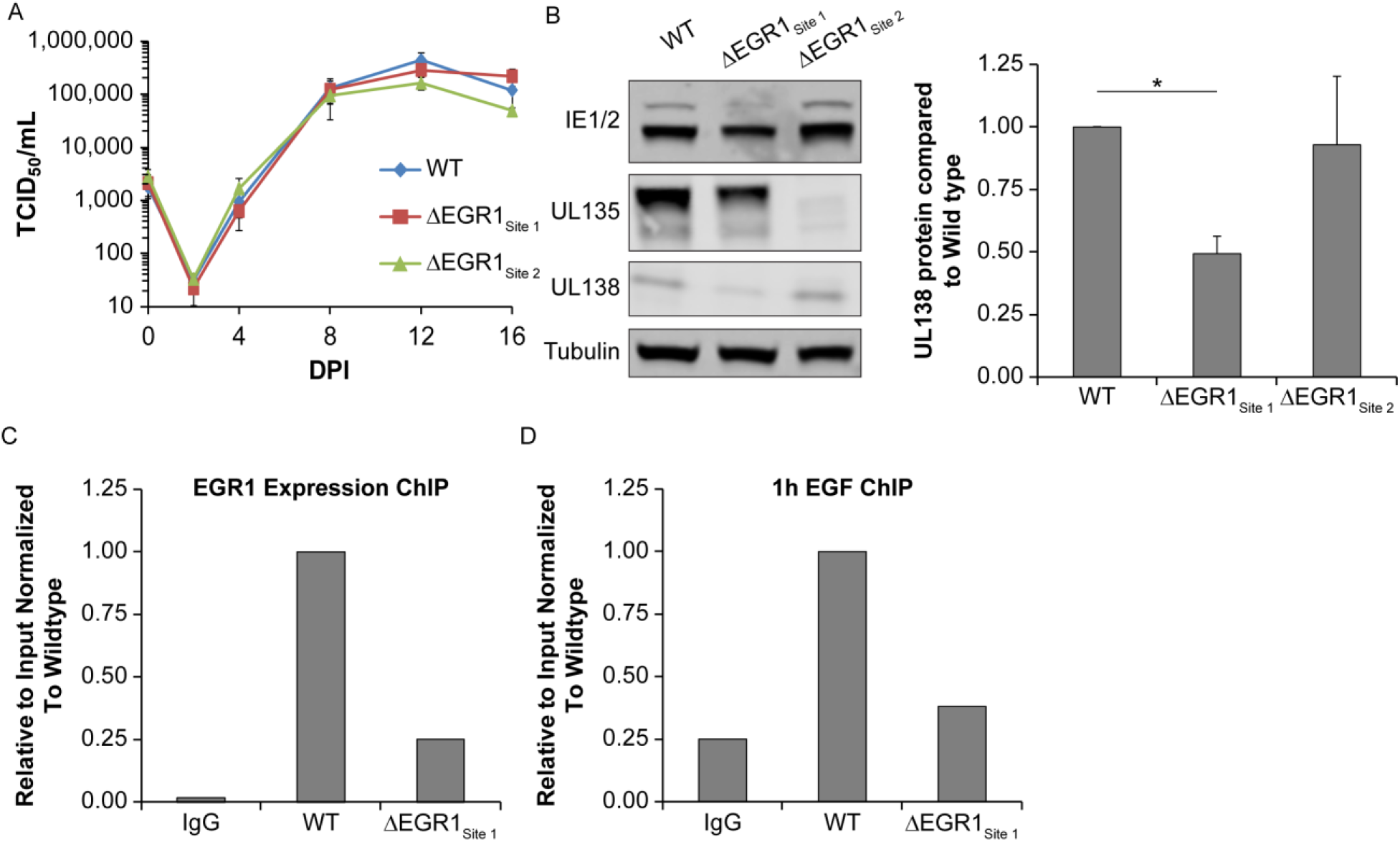
Disruption of EGR1 site 1 blocks EGR1 binding during infection. (A) Fibroblasts were infected with wild type TB40E_GFP_ or EGR1 binding mutant viruses, ΔSite 1 or ΔSite 2 (MOI=0.02). Cells and media were collected from 0-16 dpi and virus titers measured by TCID_50_. The average of 3 independent replicate experiments is shown. (B) Fibroblasts were infected wild type or EGR1 binding mutant viruses. Samples were lysed at 48 hpi, separated by SDS-PAGE and blotted for ms α-IE1/2, rb α-UL135, and rb α-UL138, and ms α-tubulin. UL138 protein levels were quantified and each mutant was normalized to WT over 3 independent experiments. The average value is graphed with error bars representing SEM and statistical significance is calculated by One-way ANOVA with Bonferroni correction (* p-value <0.05). (C) Fibroblasts were transduced with EGR1_3xFlag_ lentivirus for 24 h and then infected with wild type TB40EGFP or TB40E-ΔEGR1_Site 1_ mutant virus (MOI=1). After 48 h, samples were immunoprecipitated for either IgG or EGR1 and processed for ChIP-qPCR using SimpleChIP Enzymatic Chromatin IP Kit (Cell Signaling). (D) Fibroblasts were infected with wild type TB40E_GFP_ or TB40E-ΔEGR1_Site 1_ mutant virus (MOI=1) and transferred to serum-free media at 24 hpi. At 48 hpi, samples were pulsed with 10 nM EGF for 1h and processed for ChIP-qPCR as was done in E. For both E and F, the presence of EGR1 Site 1 sequence was calculated relative to a 2% input control (*Relative expression* = 0.02 × 2^(*CT*_2% *input*_-*CT*_*ChIP*_)^) and normalized to wild type levels.

To determine if either EGR1 binding site mutation affected *UL138* expression in the context of infection, we infected fibroblasts with TB40/E-WT, -ΔEGR1_Site 1_ or -ΔEGR1_Site 2_, and measured UL138, UL135, and IE1/IE2 protein levels at 48 hpi by immunoblot (Fig. 7B). Disruption of Site 1, but not Site 2, decreased *UL138* protein levels by 50%. While, IE protein levels were unaffected, we detected a profound loss of UL135 protein in the TB40/E-ΔEGR1_Site 2_ infection. This loss was evident in multiple independent clones of this virus. Given the importance of UL135 on reactivation (18), we moved forward with only the TB40/E-ΔEGR1_Site 1_ mutant virus.

To confirm the loss of EGR1 binding to ΔEGR1_Site 1_ in the context of infection, we transduced fibroblasts with EGR1_3xFlag_ lentivirus and infected with either WT or TB40/E-ΔEGR1_Site 1_ mutant virus. EGR1 was immunoprecipitated (SimpleChIP, Cell Signaling Technologies) and binding to site 1 was quantified by qPCR using primers flanking site 1. EGR1 precipitated 50-fold more site 1 sequence in the WT-virus infection relative to the IgG control (Fig. 7C). Infection with TB40/E- Δ EGR1_Site 1_ resulted in a 4-fold reduction in EGR1 binding relative to wild-type. We also analyzed binding of endogenous EGR1 to Site 1 in fibroblasts infected with either TB40/E-WT or TB40/E-ΔEGR1_Site 1_ and stimulated with EGF for 1h to induce EGR1 (Fig. 7D). As in Figure 7C, EGR1 binding to site 1 was increased 4-fold relative to IgG control and was reduced nearly 4-fold in TB40/E-Δ EGR1_Site 1_ infection relative to. Taken together, these data demonstrate that disruption of the EGR1 binding site 1 in the UL133/8 regions disrupts EGR1 binding and EGR1-mediate induction of *UL138* expression.

### EGR1-stimulation of UL138 is required for CMV latency

To determine if EGR1 binding is important for CMV latency we infected CD34^+^ HPCs with either TB40/E-WT or TB40/E- ΔEGR1_Site 1_ mutant virus. At 24 hpi, pure populations of infected HPCs (GFP^+^CD34^+^) were isolated and seeded into transwells above a stromal cell support for long-term culture. After 10 days of long-term culture, the cultures were split and half were mechanically lysed. We then seeded the lysate or live cells by limiting dilution in parallel onto fibroblasts. GFP^+^ wells were scored 14 days later to determine the frequency of infectious centers present at the time of lysis (pre-reactivation) or resulting from reactivation, as described for Figure 3B. Reactivation of TB40/E-WT produced a 3-fold increase in the frequency of infectious centers relative to the pre-reactivation control (Fig. 8). In contrast, TB40/E-ΔEGR1_Site 1_ infection resulted in a loss of latency and equal frequencies of infectious centers were measured prior to and following reactivation. The loss of latency with the TB40/E-ΔEGR1_Site 1_ mutant is a similar phenotype as a *UL138*_*null*_-mutant virus in CD34^+^ HPCs (18). While the cell numbers required for immunoblots of viral proteins is prohibitive, the loss of latency phenotypes are consistent with diminished expression of *UL138*. Further, these results are consistent with our finding that inhibition of EGFR, PI3K, or ERK1/2 signaling promotes the reactivation of viral replication (Fig. 3B).

**Figure 8.**
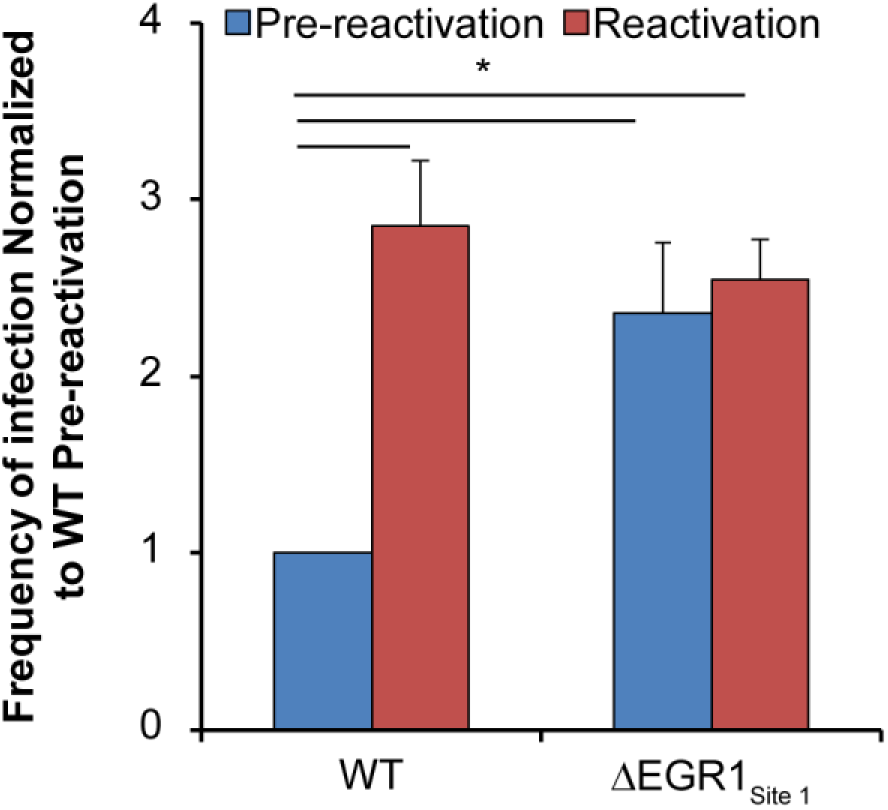
Inhibition of EGR1 binding to site 1 disrupts CMV latency. (A) CD34^+^ HPCs were infected with either wildtype TB40E_GFP_ or TB40E-ΔEGR1_Site 1_ mutant virus (MOI=2). At 24 hpi, CD34^+^/GPF^+^ cells were sorted and seeded into long-term culture. After 10 days in culture, parallel populations of either mechanically lysed cells or whole cells were plated onto fibroblast monolayers in cytokine-rich media. 14 days later, GFP+ wells were scored and frequency of infectious centers was determined by extreme limited dilution analysis (reactivation). The mechanically lysed population defines the quantity of virus present prior to reactivation (pre-reactivation). The frequency was normalized to wild type pre-reactivation and the average of three independent experiments is shown. Statistical significance was calculated by one-way ANOVA with Tukey’s correction and represented by asterisks (* p-value < 0.05).

## DISCUSSION

To regulate the establishment of latency and the reactivation of replication, all herpesviruses rely on and manipulate cellular signaling pathways to regulate their viral lifecycle. CMV targets and manipulates EGFR and its downstream pathways to achieve this goal (11, 13, 15, 19, 24, 26). Stimulation of EGFR signaling upon entry into hematopoietic cells is important to establish an environment to support latency in CD34+ HPCs (10, 16, 45). While EGFR is downregulated during the replicative cycle, targeting EGFR provides CMV with access to cellular processes involved in differentiation, proliferation, motility, immune signaling, and DNA repair (46–50). By incorporating EGFR signaling into CMV entry, latency, and reactivation of viral replication the virus maintains a firm grasp on the receptor and its signaling cascades (13). From the findings of this study, we propose a model whereby EGFR signaling through MEK/ERK and PI3K/AKT pathways drives *UL138* expression through the induction of EGR1 to suppress viral replication for latency (Fig. 9). As such, the high levels of EGR1 in CD34^+^ HPCs likely primes these cells for expression of *UL138* upon infection and the establishment of latency upon infection (21). Combined with our previous findings that *UL138* maintains EGFR surface levels (15), we propose a model by which EGFR and *UL138* form a positive feedback loop to promote and maintain CMV latency. By disrupting the feedback loop, with either chemical inhibitors or preventing EGR1 binding, the virus is unable to maintain latency and replicates. We have previously shown that UL135 opposes UL138 in targeting EGFR for turnover and reactivation (15, 19). Here, we demonstrate that CMV miR-US22 also counteracts UL138 regulation of EGR1 by targeting EGR1.

**Figure 9.**
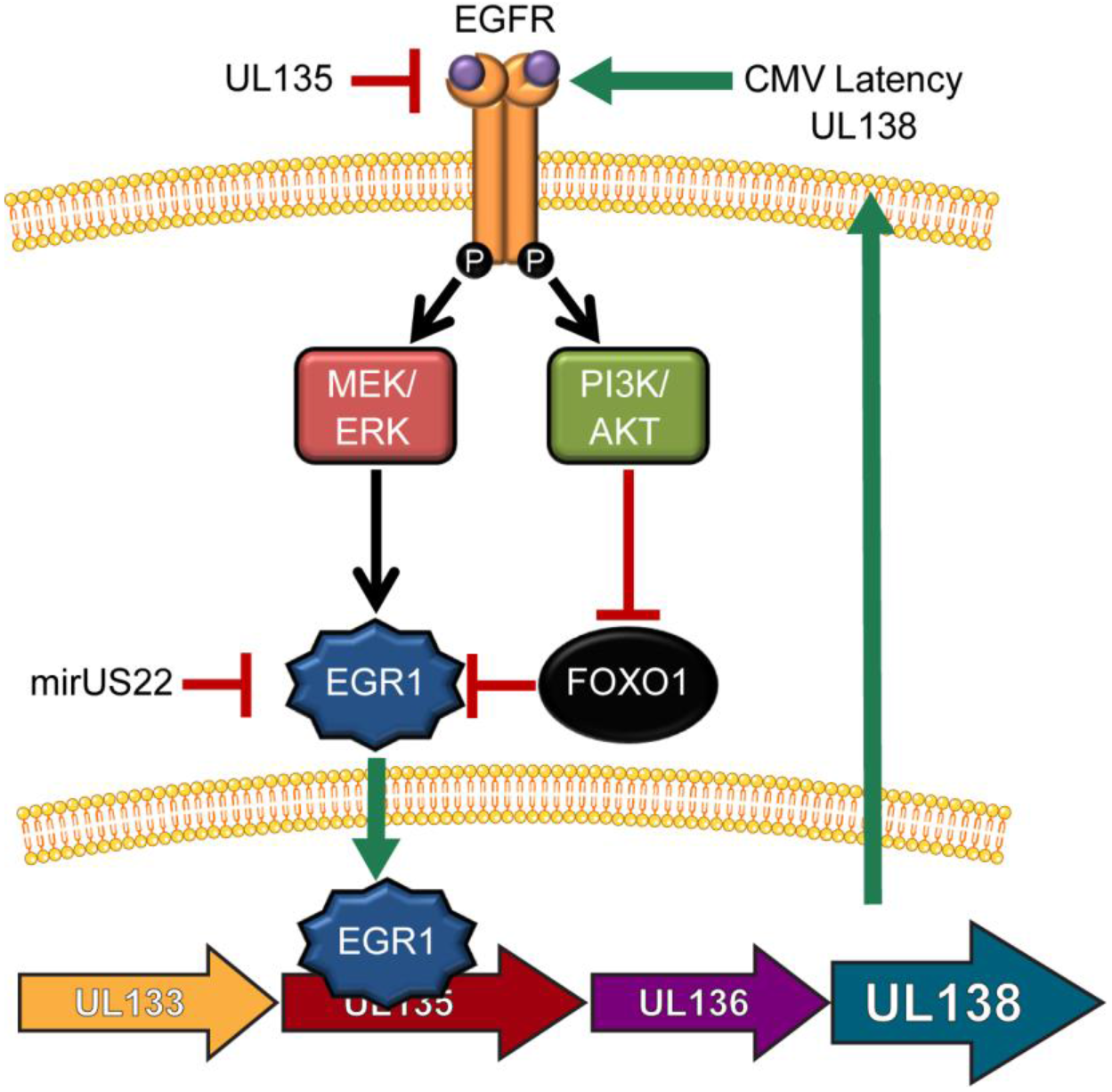
UL138 expression is regulated by a positive feedback mechanisms through EGFR signaling. Our data demonstrates that EGFR signaling promotes *UL138* expression through an upstream EGR1 binding site within a promoter in the UL133/8 locus that remains to be mapped. The EGFR signaling pathway drives the establishment of latency, at least in part, by stimulating *UL138* expression, which can function to sustain EGFR signaling. While we have previously shown that UL135 targets EGFR for turnover, miR-US22 provides an additional point of control for reactivation in targeting EGR1.

We demonstrate that EGFR activation of the MEK/ERK and PI3K/AKT signaling axes regulates gene expression of *UL138* through EGR1 binding to an unmapped element in the *UL133-UL138* locus that stimulates the expression of *UL138*, which in turn suppresses virus replication for latency. PI3K/AKT and MEK/ERK signaling pathways are common targets in herpesvirus infection and their sustained signaling maintains latency. Herpes simplex virus 1 (HSV-1) activates PI3K activity through stimulation of neural growth factor to maintain persistence (51). Inhibition of P13K stimulates HSV-1 reactivation, but full reactivation also requires c-Jun N-terminal kinase (JNK), a MAPK family member, signaling to induce histone phosphorylation on viral promoters to de-repress HSV-2 gene expression (52). Also, HSV-1 proteins VP11/12 interact with Src-family kinases, Grb2, Shc, and p85 through a tyrosine-binding motif in order to stimulate PI3K/AKT activity without growth factor stimulation (53). Additionally, HSV-1 US3 protein kinase suppresses ERK signaling to promote viral replication (54). Epstein-Barr virus (EBV) latency membrane protein-1 (LMP-1) promotes both EGFR protein levels and activation of STAT3 and ERK signaling pathways (55–57), while LMP-2A activates PI3K/AKT signaling (58, 59). Additionally, LMP-2A also promotes cellular survival through ERK activation mediating proteosomal degradation of proanoikis mediator Bim (60). Lastly, Kaposi’s sarcoma-associated herpesvirus (KSHV) latent infection promotes AKT/PI3K activation (61). However, in contrast to CMV infection, KSHV activates MEK/ERK signaling to promote its reactivation and inhibition of MEK/ERK signal suppresses ORF50 expression and KSHV reactivation (62, 63). These combined findings illustrate the significance of PI3K/AKT and MEK/ERK signaling pathways to the regulation of herpesvirus programs of latency and replication.

EGR1 induces *UL138* gene expression, but our results indicate that *UL138* is not dependent on EGR1. *UL138* is expressed from the 3’ end of a series of polycistronic transcripts encoded within the *UL133-UL138* locus and differ only in their 5’ ends (38, 40). Therefore, while EGR1 contributes to heightened *UL138* expression under specific contexts of infection in the cell, there are other mechanism that allow for constitutive or inducible *UL138* expression. One additional contributor to *UL138* expression is an IRES element that overlaps the UL136 ORF. This element is upstream of the EGR1 binding sites and stimulates downstream *UL138* gene expression from long polycistronic transcripts (38, 40). Therefore, multiple transcription and translational regulatory mechanisms control the expression of *UL138*.

The role of EGR1 in promoting *UL138* expression during CMV infection (Fig. 4 and Fig. 6) is particularly intriguing because CD34^+^ HPCs express high levels of EGR1 during maintenance in the bone marrow (21). As such, CD34^+^ cells are predisposed to promote *UL138* expression upon CMV infection to suppress viral replication. This observation is one explanation for why we have only detected UL138 protein, but not other UL133-UL138 proteins, in latently infected CD34^+^ HPCs (39). Upon differentiation of CD34^+^ HPCs EGR1 expression is abolished (21), a requirement for the differentiation and migration of stem cells out of the bone marrow. Therefore, decreased level of UL138 as a result of diminished EGR1 combined with myeloid differentiation, which would predispose the cells towards reactivation. By contrast, EGR1 levels are low in sites of productive replication, such as fibroblasts. CMV-mediated suppression of EGFR, MEK/ERK and PI3K/AKT signaling would result in the further suppression of EGR1 expression for replication (Fig. 2 and 5C). Additionally, the CMV miroRNA, miR-US22, targets EGR1 messages and is necessary for the reactivation from latency(41). In collaboration with the Nelson group, we show that miR-US22 targeting of EGR1 reduces *UL138* expression. Finally, CMV replication in fibroblasts stimulates WT1 expression (24), which competes antagonistically for EGR1 targets, including EGFR (42, 64), and indicates another mechanism by which the virus antagonizes EGFR/EGR1 signaling for productive infection. Taken together, our results demonstrate that EGR1 is an important target in CMV infection and its expression is targeted through multiple mechanisms to promote viral replication.

EGR1 regulates viral gene expression in the context of other herpesvirus infections. In HSV-1, EGR1 binding sites are located within the introns for both ICP22 and ICP4 (65). In contrast to our findings, EGR1 inhibits both ICP4 and ICP22 by blocking SP1 binding sites and recruiting the co-repressor Nab2 (65). The authors predict that the inhibition of both of these immediate early genes would promote HSV-1 gene silencing for the establishment of latency. In further contrast to CMV, during KSHV infection glycoprotein B suppresses ERK1/2 signaling to decrease expression of ORF50 to promote the establishment of latency(63). Similar to KHSV, EBV transactivator ZTA upregulates EGR1 by both interacting with its promoter and by increasing ERK signaling in order to promote viral reactivation(66). While the mechanism by which each herpesvirus utilizes EGR1 to control viral latency and replication differs, it is clear that manipulating EGR1 is common feature. These differences may reflect the unique cell types where each herpesvirus establishes latency.

Herpesviruses manipulate multiple signaling pathways to control viral latency and to promote viral replication, and understanding the complex interplay between these signaling pathways and the virus is necessary to fully appreciate how these viruses persist. We have shown that viral manipulation of host signaling impacts the control of viral transcription. Putative EGR1 binding sites exist throughout the CMV genome and, because EGR1 can either promote or repress gene expression (64, 67), it will be important to understand how EGR1 binding to promoters across the genome impact latent and replicative states of infection.

## MATERIALS AND METHODS

### Cells

MRC-5 lung fibroblasts (ATCC), HEK293T/17 cells (ATCC), Sl/Sl stromal cells (Stem Cell Technology), M2-10B4 stromal cells (Stem Cell Technology), and CD34^+^ HPCs were maintained as previously described (18). Human CD34^+^ HPCs were isolated from de-identified medical waste following bone marrow isolations from healthy donors for clinical procedures at the Banner-University Medical Center at the University of Arizona. Latency assays were performed as previously described (15, 18).

### Viruses

Bacterial artificial chromosome (BAC) stocks of TB40/E WT virus expressing GFP from a SV40-promoter was provided as gift from the Szinger lab (22). EGR1 binding mutant viruses were created by two-step, positive-negative selection approach with galK substitution as was previously described (18, 68). Both the TB40/E *UL133/8*_Null_ galK intermediate and pGEM-T *UL133-UL138* shuttle vector, referred to as *UL133/8* plasmid in the results, were created previously and described in Umashankar et al. 2014 (18). EGR1 binding sites were mutated by Phusion PCR mutagenesis using flanking PCR primers to each region with mutations incorporated into the corresponding forward and reverse primers (Table 1). pGEM-T *UL133-UL138* plasmids removing EGR1 site 1 (ΔSite 1), EGR1 site 2 (ΔSite 2), or both EGR1 sites (ΔSite 1+2) were created and stocks were maintained in DH10B bacteria glycerol stocks. Inserts for BAC recombineering were gel purified from pGEM-T ΔSite 1 and ΔSite 2 plasmids digested with EcoRI. Inserts were electroporated into SW102 *E. coli* containing the TB40/E UL133/8Null galK intermediate as previously described (69). BAC integrity was confirmed by comparing EcoRV digestion of the EGR1 binding site mutant BACs to wild type TB40/E BAC digest. Mutations of EGR1 binding sites in TB40/E-ΔEGR1_Site 1_ and TB40/E-ΔEGR1_Site 2_ were confirmed by Sanger sequencing. Loss of EGR1 binding in TB40/E-ΔEGR1_Site 1_ was confirmed by ChIP-qPCR, described below. TB40/E_GFP_ΔmiR-US22 was created as described in Mikell et al. (41).

**Table 1.**
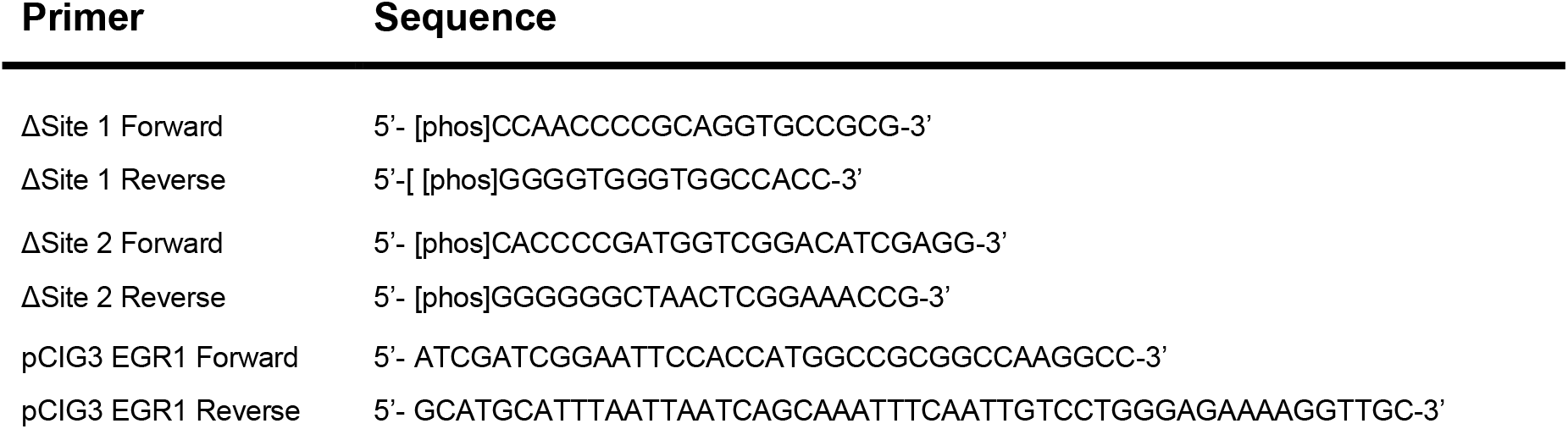
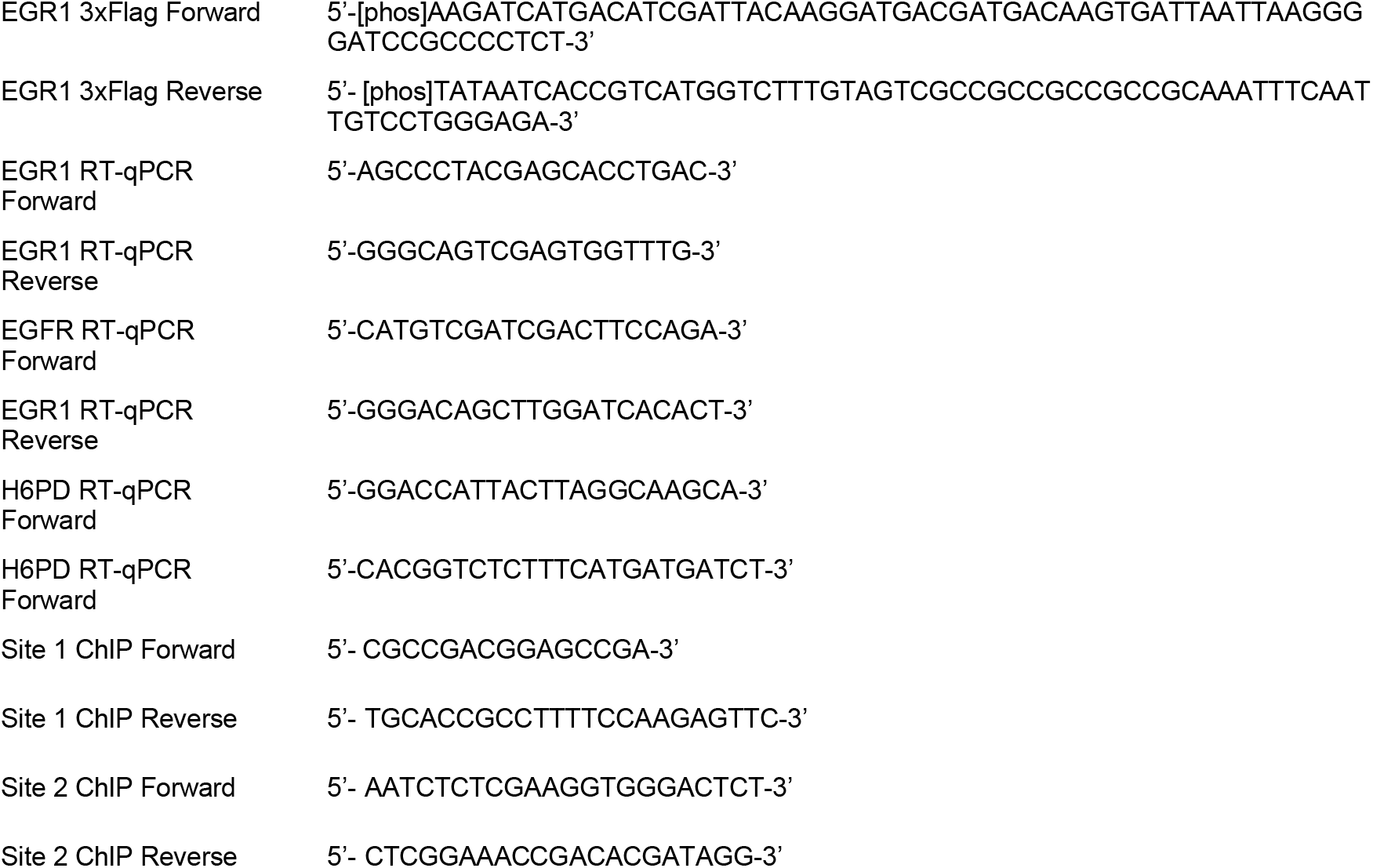
Primer sequences.

### Plasmids and Lentiviruses

pDONR221 containing EGR1 cDNA was purchased from DNASU (Arizona State University; Phoenix, Az). EGR1 was PCR amplified from the pDONR221 plasmid with pCIG3 EGR1 forward and reverse primes and inserted into pCIG3 plasmid at PacI and EcoRI digestion sites. Addition of a 3xFlag epitope tag was done by Phusion PCR mutagenesis EGR1 3xFlag Forward and Reverse primers EGR1_3xFlag_ Forward and Reverse sequences. All plasmid inserts were verified through Sanger sequencing and maintained in DH10B glycerol stocks. EGR1_3xFlag_ lentivirus was created by cotransfecting pCIG3 EGR1_3xFlag_, pMD2.G, and psPAX2 (Addgene #12259 and 12260; Trono Lab) into HEK293T/17 cells with polyethylenimine (Polysciences) and collected supernatants at 48 and 72h post transfection. Plasmid transfections were carried out in HEK293T/17 cells using PEI at 1 μg of DNA to 3 μg PEI. Plasmid encoding shRNA of EGR1 was described in Mikell et al. (41).

### Flow Cytometry

MRC-5 fibroblasts were infected with 1 MOI of TB40/E_GFP_ virus for 0-72 hpi. Cells were lifted off the plates, fixed in 2% Formaldehyde in PBS for 30 min, and washed with excess PBS. Cells were then stained with Brilliant Violent 421-conjugated ms α-EGFR (Biolegend; Table 2). Samples were gated for intact GFP^+^ cells and geometric mean of fluorescence intensity (geoMFI) was measured using a BD LSRII equipped with FACSDiva Software (BD Bioscience Immunocytometry Systems) and FlowJo software.

**Table 2.** Antibody description and Sources.

| Antibody | Species | Source | Concentration |
| --- | --- | --- | --- |
| Alexa Fluor 546 anti rabbit | goat | Molecular Probes | IF: 1:7,000 |
| Alexa Fluor 647 anti mouse | goat | Molecular Probes | IF: 1:7,000 |
| Brilliant Violet 421 EGFR | mouse | BioLegend | Flow 5 μL/ $1 \times 10^6$ cells |
| Dylight 700 conjugated anti mouse | goat | Pierce | Western: 1:12,000 |
| Dylight 800 conjugate anti rabbit | goat | Pierce | Western: 1:12000 |
| EGFR (D38B1) | rabbit | Cell Signaling | Western 1:10,00; IF 1:50 |
| EGR1 (44D5) | Rabbit | Cell Signaling | Western 1:1,000;<br>ChIP 10 μL/ $4 \times 10^6$ cells |
| EGR1 | rabbit | Bethyl | Western 1:1,000 |
| Flag (D655B) | rabbit | Cell Signaling | Western 1:1,000 |
| GAPDH (6C5) | mouse | Abcam | Western 1:15,000 |
| HRP anti-mouse | goat | Jackson ImmunoResearch | Western: 1:5000 |
| HRP anti-rabbit | goat | Jackson ImmunoResearch | Western: 1:5000 |
| IE1 (8B1.2) | mouse | Milipore Sigma | Western: 1:40,000 |
| IE1/2 (3H4) | mouse | Tom Shenk; Princeton University | Western 1:1,000 |
| IE2 | mouse | Tom Shenk; Princeton University | IF 1:50 |
| IgG, Normal | rabbit | Cell Signaling | ChIP 2 μg/ $4 \times 10^6$ cells |
| Histone H3 (D2B12) | rabbit | Cell Signaling | ChIP 10 μL/ $4 \times 10^6$ cells |
| PE conjugated CD34 | mouse | BD Biosciences | Flow 20 μL/ $1 \times 10^6$ cells |
| phospho-AKT (S473; D9E) | rabbit | Cell Signaling | Western 1:2000 |
| phospho-EGFR (Tyr1068; D7A5) | rabbit | Cell Signaling | Western 1:1000 |
| phospho-p44/42 MAPK (ERK1/2) (Thr202/Tyr204) (D13.14.4E) | rabbit | Cell Signaling | Western 1:2000 |
| phospho-MEK1/ (S217/221; 41G9) | mouse | Cell Signaling | Western 1:2000 |
| UL135 | rabbit | Open Biosystems <sup>a</sup> | Western 2 μg/mL |
| UL138 | rabbit | Open Biosystems <sup>a</sup> | Western 2 μg/mL |
| α-Tubulin (DM1A) | mouse | Sigma | Western 1:10000 |

<sup>a</sup> Custom Ordered antibody

### Immunoblotting

Lysates were separated by electrophoresis on precast 12% Tris-Bis SDS-PAGE gel (Genscript) or 4-20% precast gels (BioRad). Gels were transferred onto Immobilon-P PVDF membrane (EMD Millipore). Antibodies were incubated in with blocking solution as well. After antibody staining, blots were incubated with fluorescent secondary antibodies and imaged and quantitated using a Li-Cor Odyssey CLx infrared scanner with Image Studio software. Antibodies and sources are defined in Table 2. US22 experiments were developed using chemiluminescence with film and quantified with Image J software.

### RT-qPCR

Cells were infected with 1 MOI of TB40/E_GFP_ and RNA was isolated using Quick-DNA/RNA miniprep kit (Zymo Research) from 0-72 hpi. RNA was reverse transcribed into cDNA using Transcriptor First Strand cDNA Synthesis Kit (Roche). cDNA for EGR1, EGFR, and H6PD was quantified using LightCycler SYBR Mix kit (Roche) and corresponding primers (Table 1). Assays performed on Light Cycler 480 and corresponding software. ΔCT for EGR1 and EGFR were calculated by Pfaffl method normalized to H6PD (70).

### Immunofluorescence

Samples were processed as previously described and stained with antibodies (Table 2; (71)). All images were obtained using a DeltaVision RT inverted Deconvolution microscope. Representative single plane images were adjusted for brightness and contrast.

### EGF Pulse

Uninfected or cells infected with wild type or mutant TB40/E_GFP_ virus were washed two time with PBS and serum starved in serum-free media for 24h prior to EGF stimulation. After serum starvation, cells were washed with ice cold PBS and incubated on ice for 30 min. Cells were then incubated on ice with serum free media containing 10 nM EGF (Gold Biotechnology) for 30 min, except for no EGF control. Cells were then washed with ice cold PBS. 37°C serum free media was added and samples were incubated at 37°C for 15min to 24h, depending on experiment. Samples were then collected for immunoblotting or chromatin immunoprecipitation, depending of experiment.

### siRNA knockdown

HEK293T cells, seeded into 12-well plates the day before, were co-transfected with the indicated 400ng pGEMT plasmid and 600ng pSiren plasmid per well using Lipofectamine 2000 (Invitrogen). 24 h later, the cells were serum starved overnight in 0% FBS DMEM and then treated with 50 ng/mL EGF (Cell Guidance Systems) for 1 hour. Cells were harvested in protein lysis buffer (50mM Tris-HCl pH 8.0, 150mM NaCl, 1% NP-40, and protease inhibitors). The experiment was performed in duplicate.

### Measurement of infectious virus

Confluent fibroblasts were infected with either 1 MOI or 0.02 MOI of either wild type TB40/E_GFP_ or EGR1 mutant virus (TB40/E-ΔEGR1_Site 1_ and TB40/E- ΔEGR1_Site 2_). For pathway inhibitors, the media was changed 24 hpi with media containing inhibitor and incubated for 8 days. For EGR1 mutant virus studies, media was changed 24 hpi and samples were collected up to 16 dpi. Both cells and media were collected and then total virus was quantified by the TCID_50_ (18). Infectious centers were quantitated in CD34^+^ HPCs, as described previously (71). Frequency of infection centers were calculated using extreme limiting dilution analysis (72). For pathway inhibitors, CD34^+^ HPCs were treated with chemical inhibitors after sorting for CD34^+^ GFP^+^ populations. Inhibitor concentration, targets, and sources are listed in Table 3.

**Table 3.** Chemical Inhibitors.

| Inhibitor | Target | Source | Concentration |
| --- | --- | --- | --- |
| Binimetinib | MEK1/2 | LC Laboratories | 1 μM |
| SCH772984 | ERK1/2 | Selleckchem | 125 nM |
| Fludarabine | STAT1 | Selleckchem | 50 μM |
| S3I-201 | STAT3 | Selleckchem | 100 μM |
| LY294002 | PI3K | LC Laboratories | 20 μM |
| MK-2206 | AKT | Selleckchem | 1.25 μM |
| U73122 | PLCγ | Selleckchem | 4 μM |

### Next Generation sequencing analysis

Transcript data was acquired from a previous study conducted by Shu et al. (2). Briefly, they prepared mRNA libraries were prepared from CD34^+^ HPCs infected with TB40/E_GFP_ at a MOI of 2 at 2 and 6 dpi. Transcripts for EGR1, EGR2, EGR3, and WT1 were normalized to fragments per kilobase per million reads (FPKM) and then normalized to EGR1 levels at 2 dpi. Data from two independent experiments using different donors were combined and graphed together.

### ChIP-qPCR

For EGR1 overexpression chromatin immunoprecipitation coupled with qPCR (ChIP-qPCR), MRC-5 fibroblasts were transduced with 1 MOI of EGR1_3xFlag_ lentivirus. Transduced cells were then infected with 1 MOI of either wild type TB40/E_GFP_ or TB40/E- ΔEGR1_Site 1_ for 48h and then processed for ChIP. In EGF pulse ChIP-qPCR, fibroblasts were infected with 1 MOI of either wild type TB40/E_GFP_ or TB40/E-ΔEGR1_Site 1_ and then serum starved at 24 hpi. At 48 hpi, samples were pulsed with 10 nM EGF for 1h. All samples were then processed for ChIP-qPCR using SimpleChIP Enzymatic Chromatin IP Kit (Cell Signaling Technologies) as per manufacturer’s recommended protocol. ChIP was carried out using rb α-EGR1, rb α-Histone H3 (positive control), and Normal rabbit IgG (negative control) and with 4 x 10^6^ infected cells per reaction (Table 1). PCR was performed with primers specific to EGR1 binding site 1 and site 2 in the *UL135* open reading frame and separated on 2% agarose gel with ethidium bromide (Table 2). qPCR was performed with LightCycler SYBR Mix kit (Roche) and Site 1 Forward and Reverse primers. Relative expression was calculated against a 2% input control (*Relative expression* = 0.02 × 2^(*CT*_2% *input*_-*CT*_*ChIP*_)^). Samples were then normalized to the relative expression of the WT EGR1 ChIP.

### Pathscan Antibody Array

MRC-5 fibroblasts were infected at an MOI of 1 with TB40/E_GFP_ virus and incubated for 48h. After 48h, samples were washed with PBS twice and either lysed to measure steady state phosphorylation or pulsed with 10 nM of EGF for 30 min and then lysed to measure phosphorylation post stimulation. Phosphorylation levels were measured using the PathScan EGFR Signaling Antibody Array Kit from Cell Signaling as per manufacturer recommended protocol using protein lysates at a concentration of 1 mg/ml. Arrays were analyzed us a LiCOR Odyssey scanner at a resolution of 42μm, high quality setting, and exposure intensity of 1. Phosphorylation levels were normalized to uninfected, no EGF treatment.

### Statistical Analysis

All statistics were calculated using GraphPad Prism version 7 software. Statistics for experiments in this study were calculated using either Student T-test or analysis of variance (ANOVA) for statistical comparison, which is indicated in the figure legends with p-values for each experiment.

## ACKNOWLEDGEMENTS

We acknowledge Dr. Shu Cheng at the University of Arizona for helpful discussion and providing transcript data for this manuscript. We acknowledge Dr. Luwanika Mlera for critical reading of the manuscript. We acknowledge Mark Curry and the Arizona Cancer Center/Arizona Research Laboratories Division of Biotechnology Cytometry Core Facility for expertise and assistance in flow cytometry and Patricia Jansma of the Molecular and Cellular Biology Imaging Facility for expertise in and assistance in fluorescent imaging. Special thanks to Terry Fox Laboratory for providing the M2-10B4 and Sl/Sl cells. We acknowledge Dr. Tom Shenk for the gift of antibodies.

This work was funded by the National Institutes of Health R01 (R01 AI079059) funded to F.G, a National Institutes of Health R01 (R01 AI21640) funded to J.N., a National Institutes of Health P01 (P01 AI127335) funded to F.G. and J.N., and a National Cancer Institute institutional T32 training grant (T32CA009213-36 2014) and American Cancer Society Post-Doctoral Research Fellowship (129842-PF-16-212-01-TBE) funded to J.B.

## SUPPORTING INFORMATION

**Figure S1.**
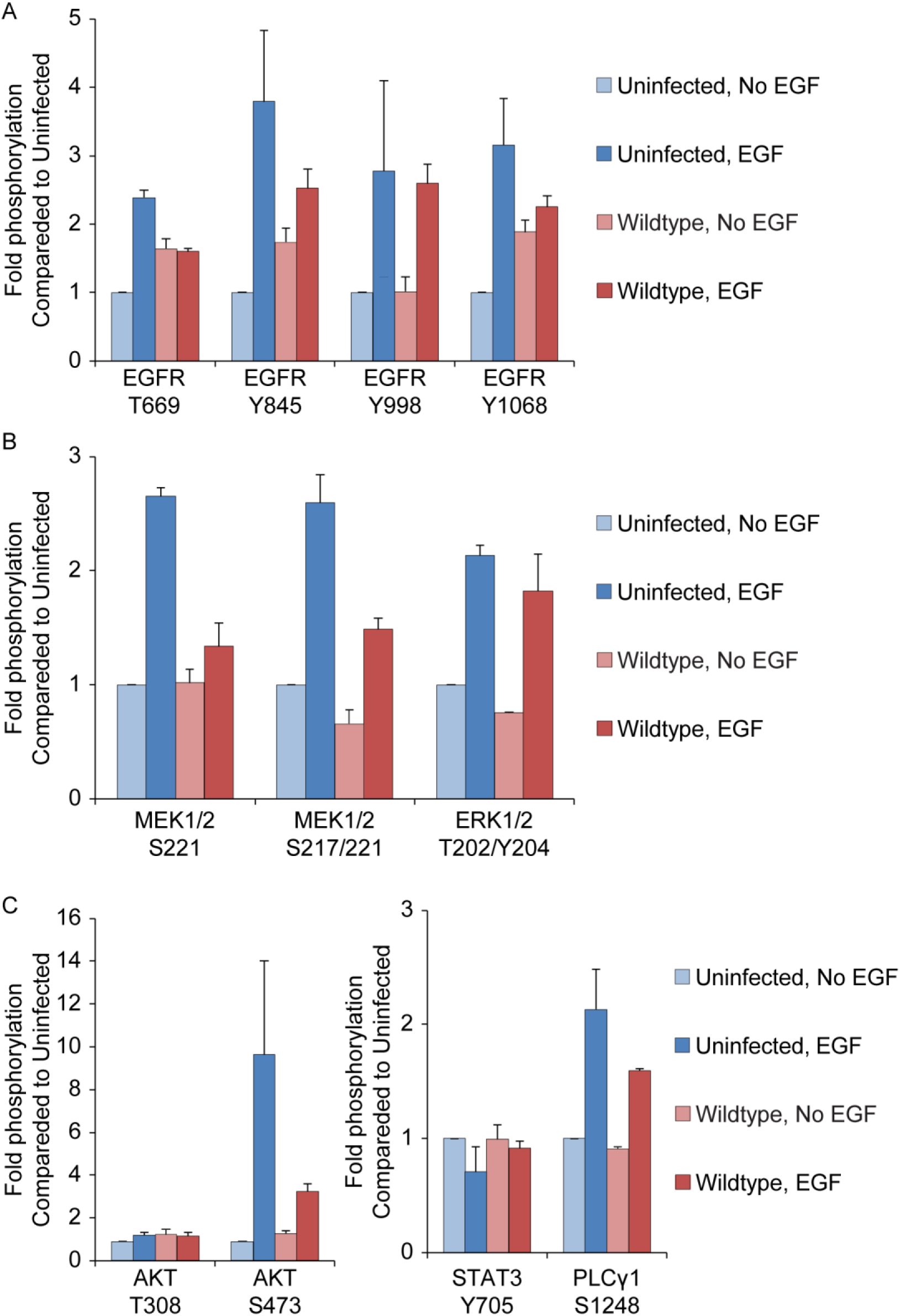
Phosphorylation screen of EGFR signaling pathways during CMV infection. Fibroblasts were infected with TB40E_GFP_ (MOI=1) for 48 h. Cells were then stimulated with 10 nM EGF for 30 min and lysed for PathScan EGFR Signaling Antibody Array Kit (Cell Signaling) analysis. Parallel unstimulated samples were lysed for comparison. Phosphorylation levels for EGFR, MEK/ERK, AKT, and STAT3 markers were normalized to uninfected, no EGF levels and graphed. Data represents two independent screens each containing two internal technical replicates. Error bars represent the range of the means from each experiment.

**Figure S2.**
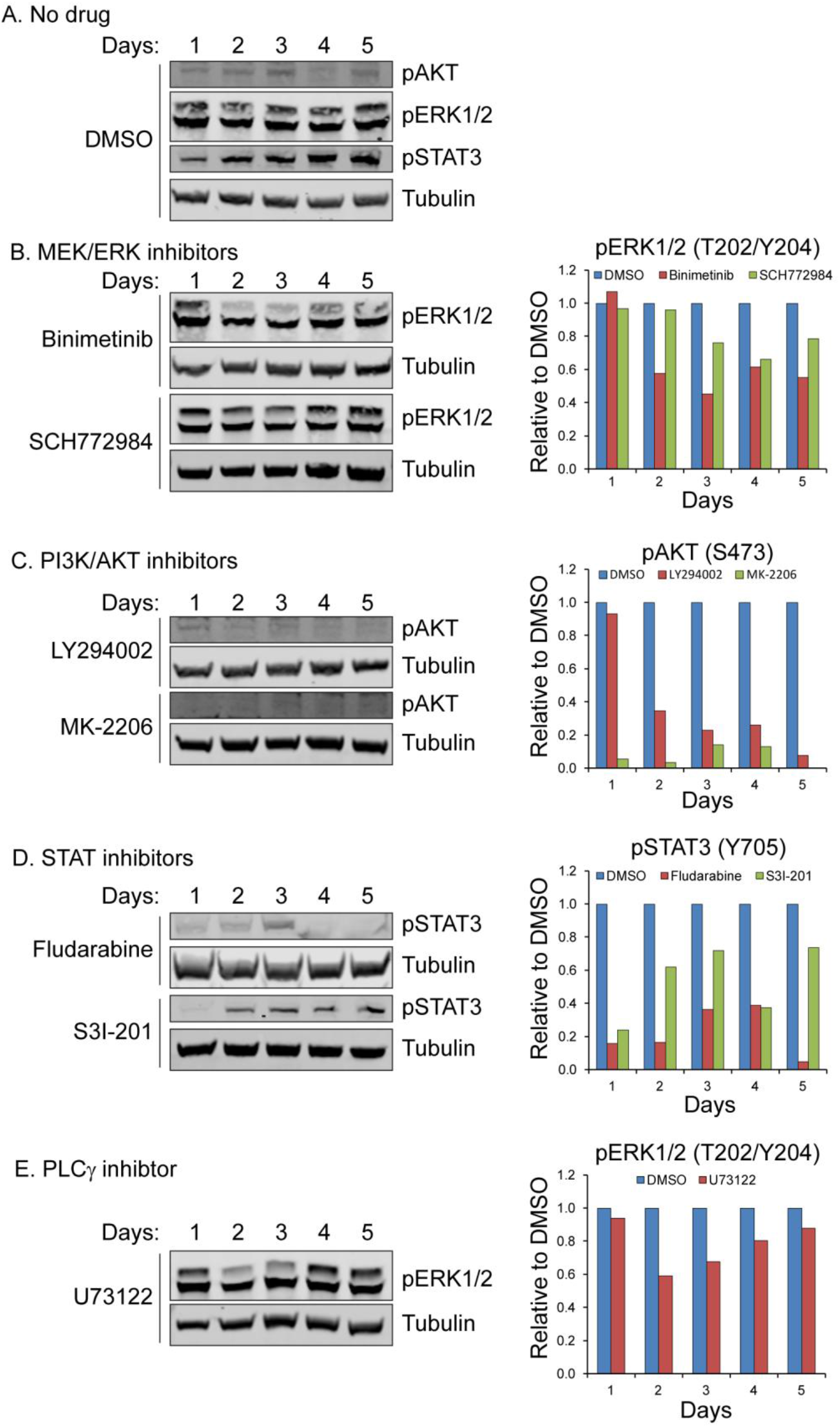
Confirmation of chemical inhibition. Fibroblasts were treated with (A) DMSO, (B) MEK/ERK inhibitors (Binimetinib; SCH772984), (C) STAT (Fludarabine; S3I-201), (D) PI3K/AKT (LY294002; MK-2206), (E) PLCγ (U73122) and lysates were isolated from 1-5 days. Samples were separated by SDS-PAGE and blotted for rb α-pAKT(S472), rb α-pERK1/2(T202/204), rb α-pSTAT3(Y705), ms α-IE1/2 antibody, and ms α-Tubulin. Inhibitor protein phosphorylation levels were normalized to DMSO controls.

**Figure S3.**
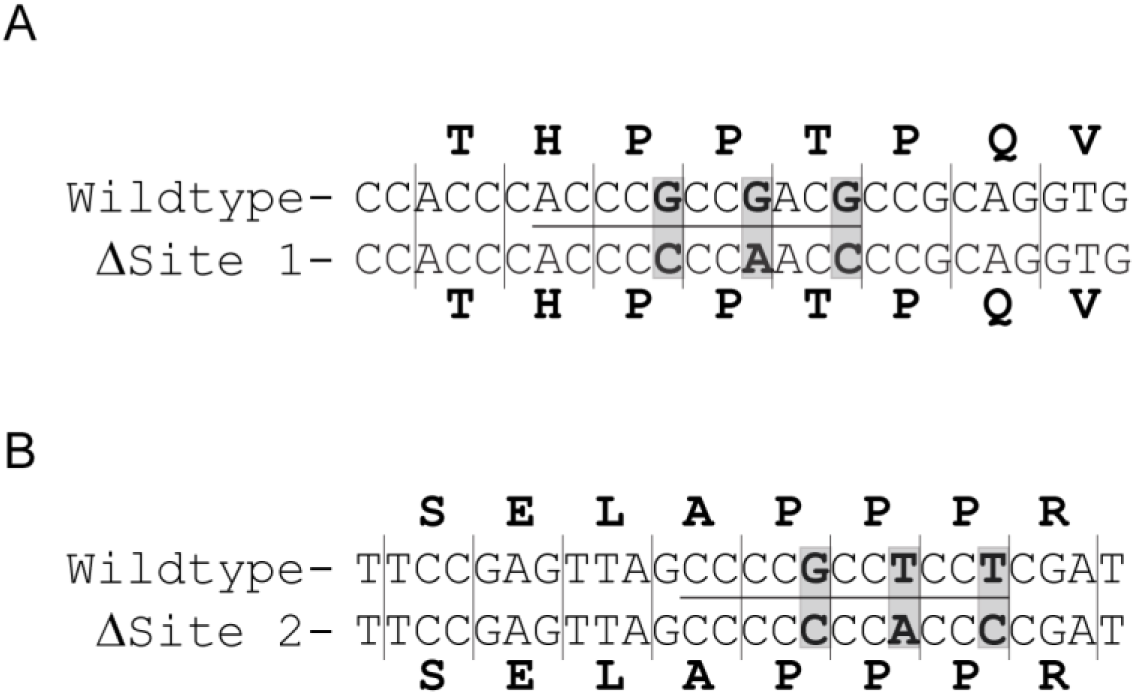
Diagram of EGR1 binding site mutation. *UL135* nucleotide sequence was altered in both a pGEM-T virus plasmid and TB40E_GFP_ bacteria artificial chromosome backbone to disrupt EGR1 binding site 1 (A) and EGR1 binding site 2 (B). Mutations were engineered into the wobble codon in order to alter the nucleotide sequence but not the amino acid sequence of UL135. Binding sequence for each site is underlined and nucleotides mutated are indicated in grey boxes and bolded text.

